# Molecular consequences of acute versus chronic CDK12 loss in prostate carcinoma nominates distinct therapeutic strategies

**DOI:** 10.1101/2024.07.16.603734

**Authors:** Sander Frank, Thomas Persse, Ilsa Coleman, Armand Bankhead, Dapei Li, Navonil De-Sarkar, Divin Wilson, Dmytro Rudoy, Manasvita Vashisth, Patty Galipeau, Michael Yang, Brian Hanratty, Ruth Dumpit, Colm Morrissey, Eva Corey, R. Bruce Montgomery, Michael C. Haffner, Colin C. Pritchard, Valeri Vasioukhin, Gavin Ha, Peter S. Nelson

## Abstract

Genomic loss of the transcriptional kinase *CDK12* occurs in ∼6% of metastatic castration-resistant prostate cancers (mCRPC) and correlates with poor patient outcomes. Prior studies demonstrate that acute CDK12 loss confers a homologous recombination (HR) deficiency (HRd) phenotype via premature intronic polyadenylation (IPA) of key HR pathway genes, including *ATM.* However, mCRPC patients have not demonstrated benefit from therapies that exploit HRd such as inhibitors of polyADP ribose polymerase (PARP). Based on this discordance, we sought to test the hypothesis that an HRd phenotype is primarily a consequence of acute *CDK12* loss and the effect is greatly diminished in prostate cancers adapted to *CDK12* loss. Analyses of whole genome sequences (WGS) and RNA sequences (RNAseq) of human mCRPCs determined that tumors with biallelic *CDK12* alterations (*CDK12^BAL^*) lack genomic scar signatures indicative of HRd, despite carrying bi-allelic loss and the appearance of the hallmark tandem-duplicator phenotype (TDP). Experiments confirmed that acute CDK12 inhibition resulted in aberrant polyadenylation and downregulation of long genes (including *BRCA1* and *BRCA2*) but such effects were modest or absent in tumors adapted to chronic *CDK12^BAL^*. One key exception was *ATM*, which did retain transcript shortening and reduced protein expression in the adapted *CDK12^BAL^* models. However, *CDK12^BAL^*cells demonstrated intact HR as measured by RAD51 foci formation following irradiation. *CDK12^BAL^* cells showed a vulnerability to targeting of CDK13 by sgRNA or CDK12/13 inhibitors and *in vivo* treatment of prostate cancer xenograft lines showed that tumors with *CDK12^BAL^*responded to the CDK12/13 inhibitor SR4835, while CDK12-intact lines did not. Collectively, these studies show that aberrant polyadenylation and long HR gene downregulation is primarily a consequence of acute CDK12 deficiency, which is largely compensated for in cells that have adapted to CDK12 loss. These results provide an explanation for why PARPi monotherapy has thus far failed to consistently benefit patients with CDK12 alterations, though alternate therapies that target CDK13 or transcription are candidates for future research and testing.

## INTRODUCTION

Large scale genomic analyses of localized and metastatic prostate cancers (PC) have identified a large spectrum of recurrent somatic alterations that involve the activation of oncogenic signaling pathways or the inactivation of tumor suppressor processes (1–5). For example, the androgen receptor (AR) serves as a key therapeutic target for most metastatic PCs, and recurrent somatic events that drive persistent AR activity promote treatment resistance and the emergence of metastatic castration resistant PC (mCRPC). A subset of other recurrent genomic alterations observed in mCRPC confer differential sensitivity to specific treatments: notably, mutations in genes involved in homology directed DNA repair (HR), such as *BRCA2*, confer responses to poly ADP ribose polymerase (PARP) inhibitors, and mutations in genes involved in DNA mismatch repair such as *MSH2* and *MSH6* associate with exceptional responses to immune checkpoint blockade (6–8). In addition to other frequent genomic aberrations that include gene fusions involving *TMPRSS2* and *ERG*, mutations in *TP53*, and loss of *PTEN*, there is a ‘long tail’ of genes altered in 3-10% of mCRPCs (5). Though by definition, genes comprising the ‘long tail’ involve smaller subsets of patients, the high prevalence of PC in the population means that events occurring at low frequency will still affect thousands of individuals. The classification of tumors with both common and rarer genomic alterations may aid prognosis and prioritize the allocation of treatments.

The gene encoding c*yclin dependent kinase 12* (*CDK12)* is functionally compromised by bi-allelic mutation or copy loss in about 5% of mCRPC cases (and in 1-2% of primary prostate cancers)(1, 9, 10). *CDK12* is also lost in 3-4% of ovarian cancers (10, 11). CDK12 is a transcription-associated kinase that pairs with Cyclin K (*CCNK*) to form an active complex that phosphorylates the RNA Polymerase II (RNAP II) C-terminal tail (12–14). A germline *Cdk12* knockout is embryonic lethal in mice (15). CDK12 has been reported to regulate mRNA splicing, suppress upstream intronic polyadenylation sites (IPAs), and maintain transcriptional elongation, especially for very large genes (16–20). Several genes involved in DNA repair, especially members of the HR pathway, are large (i.e. >50kb) and have been reported to be selectively downregulated in *CDK12* loss models (15, 16, 18, 21). This has led to a proposed outcome whereby *CDK12* loss in patient tumors may phenocopy HR deficiency (HRd). This is of notable clinical relevance as HRd tumors respond well to genotoxic platinum chemotherapy and PARP inhibitors (PARPi), which have proven to be effective across a range of cancers with underlying HR gene mutations, including mCRPC (6, 22–25).

Prior studies have evaluated the consequences of *CDK12* deficiency in a variety of experimental models, though nearly always under acute loss conditions including: protein degraders (12, 26, 27), *CDK12* knockdown or genetic knockout (15, 16), or treatment with small molecule CDK12/13 inhibitors (18, 21, 28). Acute CDK12 loss in cell models results in the down-regulation of HR associated genes, suppression of HR-mediated DNA repair, and synthetic lethality with PARP1/2 inhibitors (15, 21, 29). However, an HRd phenotype resulting from *CDK12* loss has not been confirmed in assessments of patient tumors or clinically with therapeutics that exploit HRd, and there are conflicting reports on the presence of HRd-associated genomic ‘scars’ in *CDK12* mutant cancers (9, 30–33). Crucially, mCRPC patients with *CDK12*-alterations have shown poor responses to PARPi, despite the fact that *CDK12* mutation is a labeled indication for two approved PARP inhibitors: olaparib (34–36) and talazoparib (37). A notable deficiency in the field has been the lack of models with a stable *CDK12^-/-^*genotype, as *CDK12* loss is poorly tolerated and attempts to generate long-term stable cell lines models have failed with the exception of an engineered *CDK12^-/-^*ovarian cancer line, which notably does not exhibit cisplatin or PARPi sensitivity (38–40). The objectives of this study were to address the molecular and phenotypic discordance between the preclinical studies associating *CDK12* loss with HRd, and *in vivo* human pathobiology, and identify vulnerabilities in PCs with *CDK12* loss that have potential applications for clinical management.

## RESULTS

### Identification of genomic characteristics that associate with *CDK12* loss in prostate cancer

To ascertain genomic and phenotypic alterations that associate with biallelic *CDK12* loss (*CDK12^BAL^*) in PC, we analyzed several large datasets where deep molecular assessments of tumors included analyses of genomic alterations by whole exome sequencing (WES) or whole genome sequencing (WGS), and metrics of gene expression by RNAseq. Four datasets with these criteria were evaluated: the SU2C/PCF International study of mCRPC comprising 442 tumors (SU2C-I), the SU2C/PCF US West Coast study of mCRPC comprising 101 tumors (SU2C-WC), the Hartwig Foundation study of metastatic PC comprising 168 tumors (HMF), and the University of Washington Autopsy study of mCRPC comprising 269 tumors from 127 patients (UW)(1, 3, 4, 41). Collectively, 39 tumors from 832 patients (4.7%) with at least 20% tumor cellularity were classified as *CDK12^BAL^*. These grouped by: 1 with biallelic copy loss (2.5**%**), 15 with single copy loss with a pathogenic second allele mutation (38.5**%**), and 23 (31%) with biallelic pathogenic mutation (**Fig 1a, Table S1**). Of the pathogenic mutations, 31 (31%) were localized to the kinase domain (**Fig 1b**). In addition, monoallelic *CDK12* pathogenic genomic alterations were identified in 92 tumors (11%) (**Fig 1a**).

**Figure 1.**
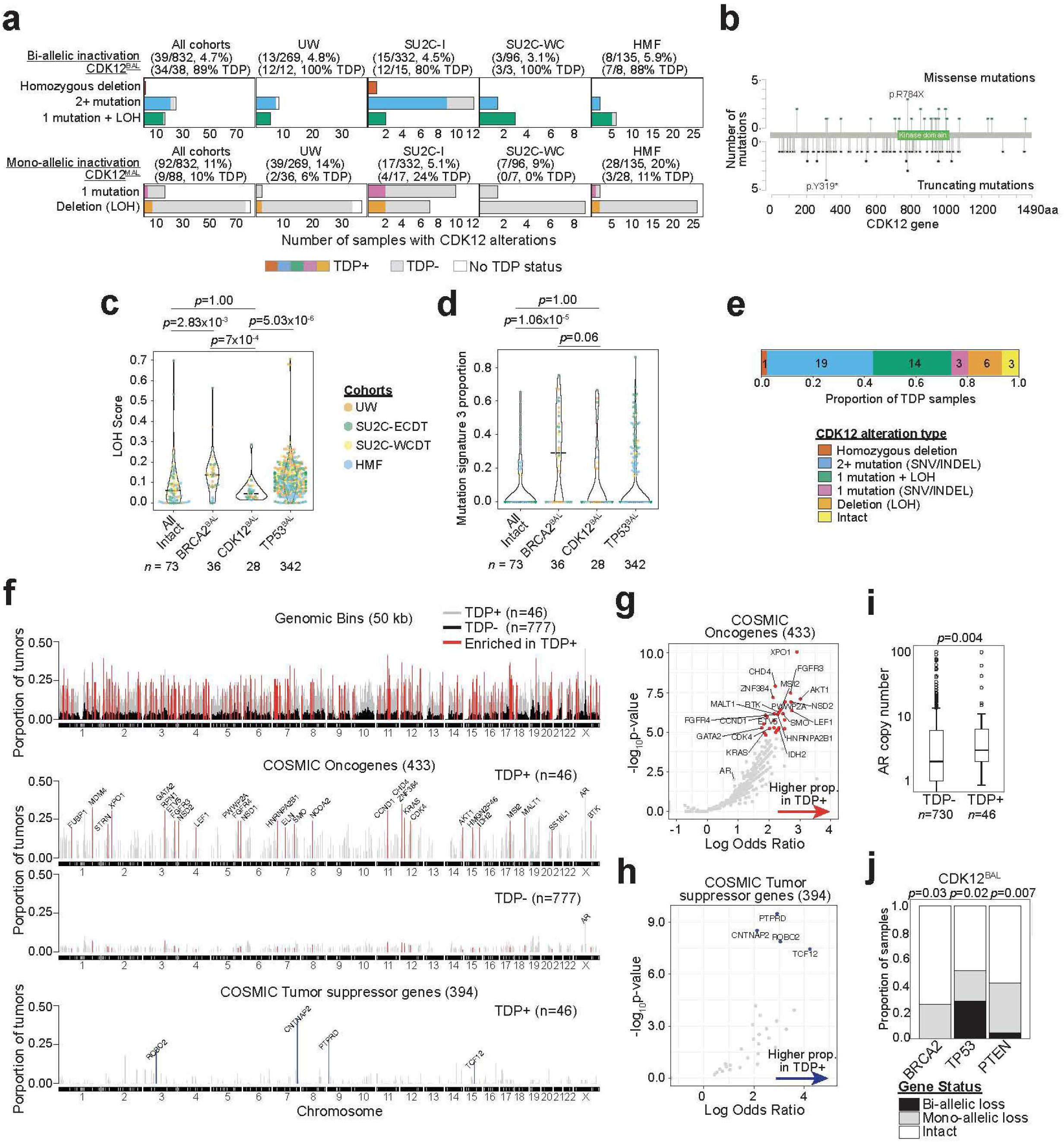
Patterns of CDK12 genomic alterations and tandem duplication phenotype. **(a)** The number of samples harboring a *CDK12* genomic alteration are shown for four CRPC cohorts. *CDK12* alterations that lead to bi-allelic loss (*CDK12^BAL^*): homozygous deletion, 2 or more (2+) small mutations (SNV/INDEL), or 1 mutation plus a loss of heterozygosity (LOH) event. CDK12 alterations that lead to monoallelic loss (CDK12^MAL^): 1 small mutation or hemizygous deletion leading to LOH. Deletion was determined based on copy loss relative to the tumor ploidy. The number of *CDK12*-altered samples with concomitant tandem duplicator phenotype (TDP) is shown in colors; grey indicates no TDP was observed; <u>white indicates sample was unevaluable for TDP (see Methods).</u>The total number and percentage of samples with *CDK12* alterations are shown in parentheses. Samples with less than 20% estimated tumor cellularity are excluded. UW, University of Washington Rapid Autopsy; SU2C-I, Stand-Up-2-Cancer International Dream Team; SU2C-WC, Stand-Up-2-Cancer West Coast Dream Team; HMF, Hartwig Medical Foundation. **(b)** Nonsynonymous mutations in *CDK12*. Counts of missense (green) and truncating (black) mutations are shown on top and bottom, respectively. Mutations included are from UW, SU2C-I, SU2C-WC, HMF cohorts (n=832) as well as panel based genomic testing (n=46). A total of 135 SNVs/INDELs in *CDK12* are present in 89 patient samples. **(c)** LOH Score for UW, SU2C-I, SU2C-WC, and HMF cohorts comparing tumors with biallelic loss of *BRCA2, CDK12, TP53*, or no alterations in these genes <u>(and in *BRCA1, CHD1, PALB2*)</u> for both alleles (All Intact). The LOH Score was defined as the proportion of the genome altered by copy number segments having zero minor copy number due to a deletion event but excluding aneuploidy involving whole or arm chromosome events. For each grouping, samples included are those with biallelic loss status of the specified gene only with absence of mutations from the other groups, or other recurrently mutated HR genes: *BRCA1, CHD1, PALB2* or genes of other groupings. ‘All Intact’ group consists of samples that do not have biallelic loss in any of *BRCA2, CDK12, TP53, BRCA1*, *CHD1* and *PALB2*. Wilcoxon rank-sum test p-value shown. **(d)** Mutational signature 3 (SBS3) that is associated with homologous recombination deficiency (COSMIC v3.4) is shown for UW, SU2C-I, SU2C-WC and HMF cohorts comparing between biallelic loss of *BRCA2, CDK12, TP53*, or no alterations in these genes (and in *BRCA1*, *CHD1*, *PALB2*) for both alleles (All Intact). Presence of SBS3 requires greater than 0.05 proportion of SNVs assigned. Mann-Whitney U test p-value shown. The samples included in the groups were selected based on the same criteria as in (c). **(e)** The proportion of *CDK12* alterations for 46 samples determined to exhibit a TDP across four CRPC cohorts. Definitions for biallelic and monoallelic loss of *CDK12* are same as in (**a**); intact refers to wildtype for both alleles. The number of samples harboring each *CDK12* alteration category is indicated. **(f)** Frequency of alteration by tandem duplications (TDs) in the 46 TDP cases. TDs were defined as simple duplication events that had flanking regions of lower copy number and length less than 10 mega base-pairs (see Methods). (Top) For each 50 kb genomic bins, the proportion of samples overlapping TDs are shown for TDP+ (grey) and TDP-cases (black). Regions with significant enrichment of TD overlap by χ^2^-test of independence (red, Bonferroni adjusted p < 0.001) spanning a total of 35.8 Mb across the genome. (Middle) Cancer Gene Census oncogenes and/or fusions whereby the gene, based on its start and end coordinates, is fully contained within a TD event. Genes with significant enrichment for being altered by TDs in TDP+ samples relative to TDP-samples by Fisher’s exact test (Bonferroni adjusted p-value < 0.01) are shown in red. (Bottom) Cancer Gene Census tumor suppressor genes and/or fusions whereby the gene is transected (broken) by either side of the TD boundary or both. Genes with significant enrichment for being altered by TDs in TDP+ samples relative to TDP-samples by Fisher’s exact test (Bonferroni adjusted p-value < 0.01) are shown in blue. **(g-h)** Genes enriched for being altered by TDs in TDP+ (n=46) versus TDP-(n=775) tumors. Shown are 433 COSMIC Cancer Gene Census oncogenes and fusions (**g**) and 390 tumor suppressor genes and fusions (**h**). Genes with-log_10_(p-value) > 5.5 and log odds ratio > 2 are shown in red for oncogenes and blue for tumor suppressors. Higher log odds ratio indicates that the proportion of the TD events altering a gene higher in TDP+ cases. Definitions of gene and TD event overlap same as in (**f**). **(i)** AR absolute copy number distribution between cases with *TDP+* (n=39) and *TDP-* (n=730) tumors. Mann-Whitney U test p-value is shown. **(j)** The proportions of *BRCA2, TP53*, and *PTEN* genomic alteration status (MAL, BAL or intact), for the cases with *CDK12^BAL^* (n=39). Fisher’s exact test was used to compare the proportions in *CDK12^BAL^ with CDK12^MAL^* or intact cases.

Having identified cohorts of PCs with and without *CDK12^BAL^*, we next sought to determine if *CDK12^BAL^* tumors exhibited evidence of HRd. Various mutational processes, including the loss of mechanisms that repair DNA damage through HR, produce characteristic signatures of residual structural alterations or mutations that can be classified according to the type of mutagen or compromised repair mechanism (33, 42, 43). Cancers with HRd are notable for genome instability resulting in large regions of copy loss and gain that can be scored based on metrics of loss of heterozygosity (LOH). LOH scores in PCs with *BRCA2^BAL^* were significantly greater than PCs without biallelic loss of *BRCA2*, *CDK12* or *TP53* (‘All Intact’ group)(Mann-Whitney U p=2.8e^-3^), whereas LOH scores in tumors with *CDK12*^BAL^ were not different than ‘All Intact’ tumors (p=1.0)(**Fig 1c; Table S1, Methods**).

COSMIC single base substitution (SBS) mutation signature 3 (CSig3) is associated with HRd and can be determined through WES or WGS (44). We determined that 10 of 39 tumors (26%) classified as *CDK12^BAL^* had evidence of CSig3 activity, a CSig3 proportion distribution which did not significantly differ compared to ‘All Intact’ tumors (Mann-Whitney U p=1.00)(**Fig 1d, Table S1, Methods**). In contrast, 46 of 69 (65%) PCs with *BRCA2* biallelic loss exhibited CSig3 signatures, having a trend of higher CSig3 activity than in *CDK12^BAL^* tumors (p<0.06)(**Fig 1d**).

CDK12 inactivation is documented to be associated with a tandem duplicator phenotype (TDP) classified by numerous copy gains of duplications across the genome (9, 11, 31). Of the 38 tumors with biallelic *CDK12* loss that were evaluable for a TDP, 34 exhibited a TD genome (89%) **(Fig 1a**). Of the four *CDK12^BAL^* tumors that were TDP(−), three had mutations after the key functional kinase domain (amino acids 727-1020)(**Table S1)**, which may be less likely to completely abolish protein function.

Overall, we classified 46 tumors with a TDP across all cohorts, including nine with monoallelic *CDK12* alteration and 3 with no alterations in *CDK12* (**Fig 1e, Methods**). Notably, six different TDP groups have been described, based on the size of the duplicated segment and whether the size distributions are unimodal or bimodal (45). Prior reports determined that tumors with *CDK12* loss generally categorize as Group 6 with TDs exhibiting a bimodal size of ∼230kb and ∼1.7Mb whereas tumors with *BRCA1* loss classify as Group 1 TDP with a unimodal size of ∼10kb. Of the 46 mCRPCs with a TDP, 30 (65%) classified as Group 6, 11 (37%) in Group 3 (median 2.6 Mb), and 3 (7%) as Group 2 (median

∼380kb) (**Fig S1a,b, Table S2**). One of 30 tumors that exhibited a Class 6 TDP did not have *CDK12* genomic alterations (**Fig S1b**). No tumors with biallelic *BRCA2* loss exhibited a TDP, though 8 of 31 tumors with monoallelic *BRCA2* alterations classified as a Group 6 TDP and none classified as the Group 1 TDP associated with *BRCA1* loss. Of tumors with *TP53* alterations, 23 (3.7%) classified as TDP+, though of these, 17 also had a *CDK12^BAL^* event (**Fig S1b, Table S2**). Collectively, these findings confirm prior reports detailing the unique TDP genomic structure associated with *CDK12^BAL^* which is distinct from the types of genomic scars associated with HRd.

We next sought to determine the gene composition within tandem duplications (TDs) to assess whether consistent oncogenic drivers or tumor suppressor mechanisms accompanied CDK12 inactivation. The median number of TDs per tumor with a TDP was 103 (range 61-161) and 426 (range 159-781) for WES and WGS data, respectively. We compared the frequency of copy gain by TDs from TDP+ tumors (n=46) against the frequency in tumors without a TDP (TDP-; n=777) and identified 130Mb (2,601 50kb windows) of total regions enriched in TDP+ tumors (Fisher’s exact test; Bonferroni adjusted p-value < 0.001)(**Fig 1f**, **Table S2, Methods**). Of 433 genes annotated as Cancer Gene Census oncogenes, 29 were significantly enriched as altered in TDP+ tumors, and 5 of 394 Cancer Gene Census tumor suppressor genes were transected by a TD boundary (p < 0.01) (**Fig 1f-h**, **Table S2**).

Though several notable genes with oncogenic functions were enriched in TDs, including *CCND1*, *AKT1* and *MDM4*, there were no genomic regions comprising a TD that recurred with a frequency greater than 40% across the 46 TDP+ tumors. Genes involved in HRd were not significantly altered in TDP tumors. The androgen receptor (AR) and upstream AR enhancer locus were contained within a TD in 18 and 19 of the 46 TDP+ tumors, respectively, with a significantly higher median AR copy number of 3 compared to 2 in non-TDP tumors (p=0.004, two-tailed Mann-Whitney U test) (**Fig 1i**). *CDK12^BAL^* was mutually exclusive with *BRCA2^BAL^* (p=0.03, one-tailed Fisher’s exact test). Of *CDK12^BAL^* tumors, 51% also harbored monoallelic (n=9) or biallelic (n=11) *TP53* alterations (**Fig 1j**) while *PTEN^BAL^* only occurred in 1 tumor with *CDK12^BAL^*.

While the majority of the mCRPC cohorts were comprised of single tumors from an individual patient, the UW autopsy study included three patients with *CDK12^BAL^* where multiple tumors were sampled, allowing for assessments of tumor heterogeneity with respect to *CDK12* events. In each patient, all metastatic tumors shared the same *CDK12* alteration, and all tumors exhibited a TDP with the majority of TDs shared across tumors within an individual (**Fig 2a**). These data suggest that *CDK12* alterations are early events in tumorigenesis and support the monoclonal model of metastatic PC dissemination (41, 46). However, there was evidence for continued accumulation of TDs as individual metastasis also exhibited a number of unique TDs, a subset of which encompassed oncogenic drivers such as *MYC*, *ETS1*, and *KRAS* (**Fig 2b**). For two tumors, we were able to profile gene expression by whole transcriptome RNAseq. Both tumors expressed high levels of the AR and AR program activity with highly concordant proliferation rates as determined by cell cycle progression (CCP) scores (**Fig 2c**).

**Figure 2.**
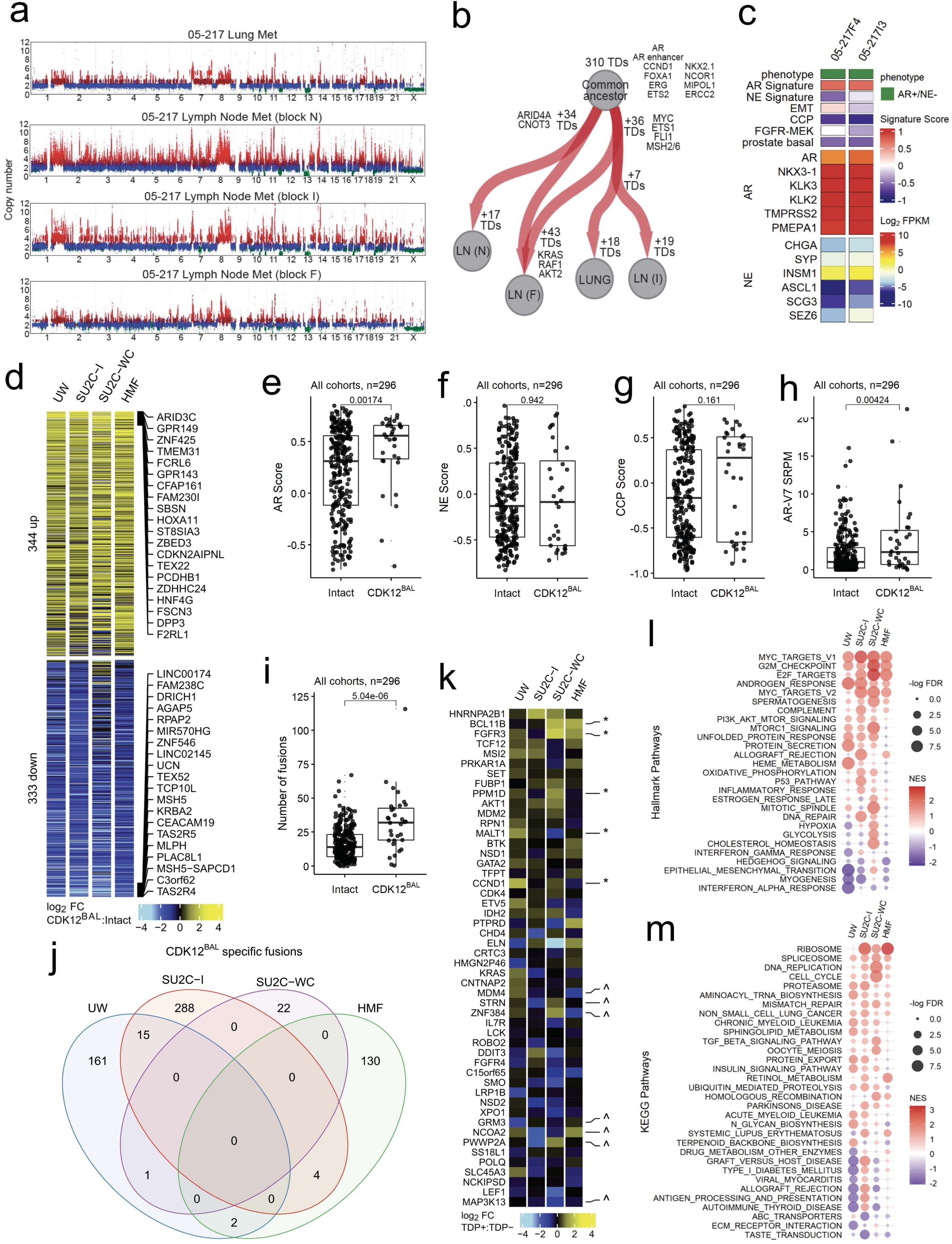
The transcriptional phenotype of mCRPCs with *CDK12* loss. **(a)** Copy number profiles for UW rapid autopsy patient 05-217 that exhibits the TDP and has *CDK12*^BAL^. Copy number alteration events of deletions (1 copy; green), copy neutral (2 copies; blue), copy gains (3 copies; dark red), and amplifications (4+ copies; red) were predicted using TITAN. Copy number values (y-axis) is shown after tumor purity and ploidy correction. **(b)** Relationship between metastatic tumors from patient 05-217 based on co-occurrence of tandem duplications (TDs). Red arrows indicate unique or shared TDs between samples. The display and orientation of the four metastases were determined by phylogenetic analysis using neighbor joining tree estimation (ape_R_package) for the presence or absence of TD events using Manhattan distance and rooted tree configuration. **(c)** Intra-patient concordance in AR active prostate cancer phenotype classification in two metastases with *CDK12^BAL^* and a tandem duplication genome. GSVA signature scores and log_2_ FPKM values are colored according to scales shown on plot. AR: androgen receptor, NE neuroendocrine, EMT: epithelial-mesenchymal transition, CCP: cell cycle progression. **(d)** RNAseq based comparison of transcript abundance differences in mCRPC tumors with and without *CDK12^BAL^* (log2 FC and p<0.05 across 2 or more cohorts). The top 20 up and down regulated gene symbols are listed **(e-h)** RNAseq based quantitation of AR and NE (neuroendocrine) pathway activity, cell cycle progression score (CCP) and levels of transcripts encoding the ARv7 splice variant comparing mCRPCs with versus without *CDK12^BAL^* (GSVA scores; Wilcoxon rank test p-values shown). **(i)** Quantitation of expressed gene fusions in mCRPCs without vs with *CDK12^BAL^*. (Wilcoxon rank test p-values shown). **(j)** Assessment of recurrence of *CDK12^BAL^*-specific expressed gene fusions within and across mCRPC cohorts. **(k)** Expression of COSMIC oncogenes and tumor suppressor genes contained within tandem duplications. * = Up-regulated or ^ down-regulated with p<0.05, FC>abs(2) in at least 1 cohort. **(l)** Hallmark Pathway enrichment in mCRPC without vs with *CDK12^BAL^* (pathways with FDR<0.05 in at least 1 cohort shown.) **(m)** KEGG Pathway enrichment in mCRPC without vs with *CDK12^BAL^* (pathways with FDR<0.05 in at least 1 cohort shown.)

### Transcriptional alterations in mCRPCs with *CDK12* Loss

While the genomic consequences of *CDK12* loss in PC and other malignancies have been established with respect to features such as tandem duplications (11, 45, 47), the assessments of phenotypic alterations in mCRPCs that accompany *CDK12* inactivation have not been evaluated extensively. As a first proxy for phenotype, we analyzed matched whole transcriptome RNAseq data from 332, 96, 135, and 269 tumors having at least 20% tumor cellularity in the SU2C-I, SU2C-WC, HMF and UW mCRPC datasets, respectively, and compared *CDK12* intact tumors against those with *CDK12*^BAL^ for differential genes, pathways, and hallmarks that reflect relevant biological characteristics of mCPRC. We removed neuroendocrine (NEPC) and AR-negative/NE-negative (DNPC) samples due to their lack of representation in the *CDK12^BAL^*group and compared the remainder to a group *CDK12* intact tumors lacking any *CDK12* or *BRCA2* alterations (n=296 tumors). Global comparisons of transcript abundance levels identified 344 genes differentially increased and 333 genes differentially decreased in *CDK12* loss vs intact tumors (log_2_ FC and p<0.05 across 2 or more cohorts)(**Fig 2d, Table S3**). There was substantial overlap in these differentially expressed genes across the four mCRPC cohorts (Bonferroni adjusted p-values<0.0001, pairwise hypergeometric tests)(**Fig S2b-c, Table S3**). We observed concordance in these datasets with previously reported gene expression alterations resulting from *CDK12* loss including the upregulation of *ARID3C*, *TBX4*, and downregulation of *TSACC* and *CDNF* in mCRPCs with *CDK12^BAL^*, along with enrichment of a CDK12-loss transcriptional signature (9)(**Fig S2a**).

mCRPCs are now recognized to exhibit subtypes classified by differentiation states reflecting androgen receptor (AR), neuroendocrine (NE), and other lineage programs. Compared to *CDK12* intact tumors, mCRPCs with *CDK12^BAL^* exhibited significantly higher AR expression and AR activity (p=0.002, Wilcoxon rank-sum test) (**Fig 2e; Fig S2d-g**). Notably, of 29 tumors classified as NEPC in the cohorts, none harbored *CDK12^BAL^* and overall NE activity scores were not different in *CDK12^BAL^* tumors (**Fig 2f**). The cell cycle progression score (CCP), a metric of cell proliferation, did not distinguish *CDK12* status (**Fig 2g**). The expression of the alternatively spliced ARv7 transcript was higher in tumors with *CDK12^BAL^* (p=0.004, Wilcoxon rank-sum test), further supporting an AR-driven phenotype (**Fig 2h**). Several pathways were reproducibly altered in *CDK12^BAL^* mCRPCs including alterations in cell cycle, androgen response, spliceosome and DNA replication (**Fig 2l,m**).

We considered several mechanisms that could explain the differential gene expression in *CDK12^BAL^* mCRPCs. We confirmed a prior study reporting high rates of gene rearrangements and fusion transcripts that associate with *CDK12^BAL^* (p=5e^-6^, Wilcoxon rank-sum test)(**Fig 2i**). However, only 22 of 623 fusion transcripts were recurrent (**Fig 2j**), and overall did not explain the differential expression of specific genes recurrently altered across studies (**Fig S2h**). In contrast, several COSMIC-defined oncogenes and tumor suppressor genes located within regions of TDs were increased at the transcript level and were recurrent across mCRPCs with *CDK12^BAL^* (**Fig 2k**). Genes involved in HR were not involved in gene fusion events and were not commonly transected by gene rearrangements or TDPs (**Fig 1h**).

A key function of CDK12 involves the regulation of gene transcription by complexing with Cyclin K to phosphorylate the C-terminal domain of RNA polymerase II which promotes transcriptional elongation and the synthesis of full-length mRNAs (14). Notably, acute loss of CDK12 in cell line models results in decreased expression of a small subset of genes that are long and comprise large numbers of exons (15). Genes with these characteristics shown to be influenced by acute CDK12 depletion include several involved in HR DNA repair such as *ATM*, *BRCA1*, *ATR*, *FANCI*, and *FANCD2* (15). To determine if the expression of long genes or those with large numbers of exons were differentially altered in PCs with *CDK12^BAL^*we analyzed the whole transcriptome RNAseq data from the four mCRPC datasets collectively and individually. We found no overall associations between differentially downregulated genes in *CDK12^BAL^*vs *CDK12*-intact tumors based on gene length when integrating tumors from all cohorts (p=0.26) (**Fig 3a**). However, other gene parameters were associated with lower transcript levels in the context of *CDK12^BAL^* including longer coding sequence length, longer transcript length, and shorter 3’UTR length (**Fig 3b, Fig S3a-c**).

**Figure 3.**
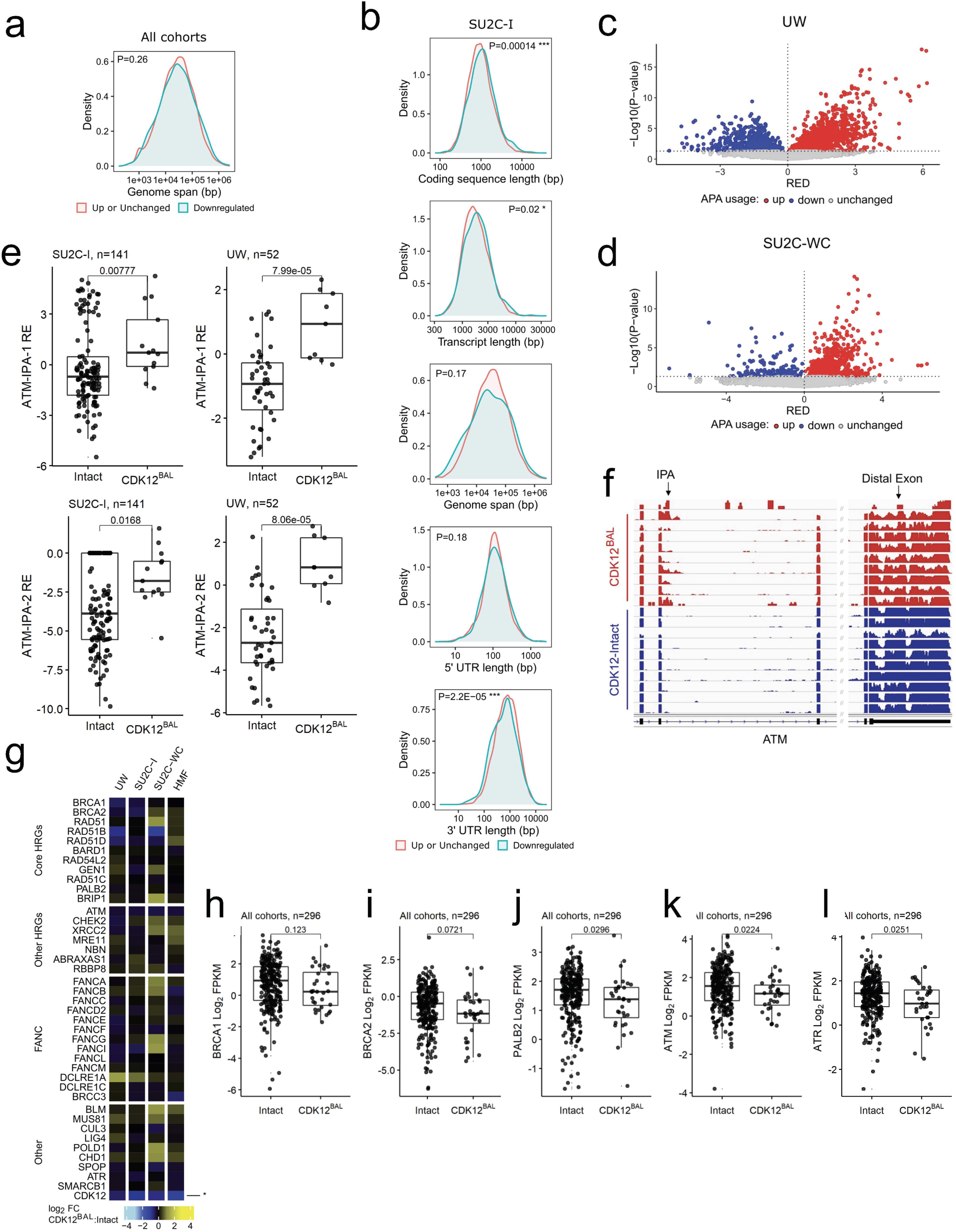
Assessment of transcript characteristics in the context of *CDK12* loss. (**a**) Analysis of transcript abundance levels determined by RNAseq comparing expression in tumors with *CDK12^BAL^* vs tumors with intact *CDK12* based on gene length (genome span). The graph represents the distribution of up-regulated or unchanged genes (red) and genes with lower transcript abundance (P<0.05; FC<-2) (blue) in *CDK12^BAL^* vs tumors with intact *CDK12*. The human gene length determination were assessed and compared using Ensemble genes with the ShinyGO tool (t-test p-value shown on plot). The y-axis shows density and the x-axis is gene length base pairs (bp). (**b**) Association of increased/unchanged or decreased (p<0.05; FC<-2) transcript abundance levels in relation to gene size features in tumors without or with *CDK12^BAL^* in the SU2C-I cohort as assessed in (a). **(c-d)** APAlyzer analysis for up– and down-regulated intronic alternative polyadenylation usage (APA) using RNA-seq from data sets comparing *CDK12^BAL^*cases vs *CDK12*-intact controls: University of Washington Autopsy study (UW), and **(d)** StandUp2Cancer (SU2C-WC). RED = relative expression difference; each point is a different APA site **(e)** Comparison of APA usage for two different IPA sites in the ATM gene in UW and SU2C-I cohortswithout vs with CDK12^BAL^ (Wilcoxon rank test p-values shown). **(f)** Transcript pile-up of reads mapping to exons demonstrating increased transcript reads corresponding to an IPA in the *ATM* gene in SU2C-I mCRPC tumors with *CDK12^BAL^* versus tumors with intact *CDK12* and diminished transcripts mapping to the distal 3’ exon. **(g)** RNAseq based transcript abundance measurements of genes involved in DNA repair, comparing mCRPCs with versus without *CDK12^BAL^*. * = Down-regulated with p<0.05, FC>abs(2) in at least 1 cohort. **(h-l)** RNAseq based transcript abundance levels for specific genes involved in the HR DNA repair pathway (Wilcoxon rank test p-values shown).

In addition to gene length-dependent effects on transcriptional elongation, acute inhibition of CDK12 activity results in premature cleavage and polyadenylation (PCPA), with the use of alternative polyadenylation sites (APA), particularly in genes with features such as large gene length, greater exon numbers, large first intron, and more intronic poly(A) sites (IPAs) (18, 48). Several genes involved in DNA damage repair fit these criteria. CDK12 has been shown to globally repress the use of intronic polyadenylation sites and the consequent expression of full-length transcripts whereas cells with CDK12 depletion exhibit elevated numbers of truncated transcript isoforms resulting from IPA usage. Analysis of previously published RNA-seq data (16) using the APAlyzer (49) tool confirmed the reported selective increase in intronic APA usage in the TCGA-PRAD primary tumors (**Fig S3d**). In mCRPC, preferential upregulation of APA sites was observed tumors with *CDK12^BAL^* in each of the mCRPC cohorts, though to variable degrees (**Fig 3c,d and Fig S3e,f**).

Several genes involved in sensing and repairing DNA damage via HR are large, and prior studies have reported downregulation of several including *ATM*, *BRCA1* and *BRCA2* in the context of acute CDK12 loss which contributes to a ‘BRCAness’ phenotype with compromised HR. We confirmed prior studies demonstrating increased use of internal polyadenylation sites in ATM and a modest reduction in transcript reads derived from the distal 3’ exon (**Fig 3e,f and Fig S2i-l**). The expression of *PALB2* and *ATR* was also modestly lower in tumors with *CDK12^BA^*^L^ (**Fig 3j,l**). However, other key HR genes were not affected as we observed no significant differential downregulation of *BRCA1* or *BRCA2* in mCRPCs with *CDK12^BAL^* nor was there evidence of APA usage in these genes (**Fig 3g-k** and data not shown).

### Differential effects of acute versus chronic *CDK12* loss on gene expression and homology directed DNA repair

We next sought an explanation for the discrepancy between the reported mechanisms of *CDK12*-loss leading to HRd and clinical observations whereby patients with *CDK12* loss exhibit poor responses to PARP inhibitors (PARPi). First, we chose to replicate acute loss conditions using approaches described in prior studies (16, 21, 26). We evaluated whether acute CDK12 inhibition caused long gene downregulation via transcript shortening, as would be expected from a role of CDK12 in suppressing APA usage. As there are no pharmacological agents that exclusively impede CDK12 activity, we used the dual CDK12/13 inhibitor SR4835 to acutely inhibit CDK12 function. Because some large DNA repair genes, including *BRCA1* and *BRCA2*, show cell cycle linked expression (50), palbociclib (Palbo) was used as a control to demonstrate the extent of mRNA decrease solely due to G1 arrest. Actinomycin D (ActD) served as a control for non-specific RNA-Pol II inhibition. Genes downregulated by SR4835 skewed longer (Students t-test p=3.2e^-4^ for LNCaP; p=3.3e^-88^ for LuCaP35_CL) than upregulated or unchanged genes, while such effects were non-significant or reversed (i.e. shorter genes downregulated) upon Palbo or ActD treatment (**Fig 4a**). This result is based on gene length (introns+exons) and not the spliced transcript length, in which case all treatments caused preferential downregulation of longer transcripts (**Fig S4a**). Though all three treatments led to some increases and decreases in APA site usage, SR4835 led to more selective enrichment for upregulated APA site usage (7,929 up, 11.0 up/down ratio) compared to ActD (3,257 up, 2.1 up/down ratio) and Palbo (1,700 up, 1.2 up/down ratio) in LNCaP with similar results in LuCaP35_CL (**Fig 4b and S4b**). After six hours of treatment, SR4835 resulted in substantial alterations in gene expression (1,128 down / 201 up in LNCaP; 2,933 down / 2,009 up in LuCaP35_CL) (**Fig. 4c**), including down regulation of multiple genes involved in the DNA repair pathways, in particular HR (13/31 in LNCaP and 22/31 in Lu-CaP_35CL) (**Fig 4d**). Pathway analysis showed that although SR4835 caused a significant decrease in the HR pathway activity by negative enrichment score (NES) (−1.1, FDR=0.9 for LNCaP, and –1.2, FDR=0.34 for LuCaP35_CL), palbociclib caused a much greater decrease (−1.9, FDR=0.001 in LNCaP, and –1.9, FDR=0.006 in LuCaP35_CL) (**Fig S4c**). In fact, approximately half (16/30 in LNCaP and 24/58 in LuCaP35_CL) of the downregulated KEGG pathways upon SR4835 treatment are also downregulated by palbociclib, leading to some difficulty in assigning which effects are due broadly to cell cycle arrest vs CDK12/13 specific effects (**Fig. S4d).** However, some genes did show SR4835-selective downregulation (e.g. *ATM*; –0.88 log2(fold), p<0.0001) and others (e.g. *RAD51D*) were downregulated more dramatically with SR4835 (−0.77, p<0.0001) than Palbo (−0.33, p=0.002) (**Fig 4e**). Thus, while the transcriptional effects from CDK12/13i treatment may be partially confounded by cell arrest effects, several key DNA repair genes do show selective downregulation.

**Figure 4.**
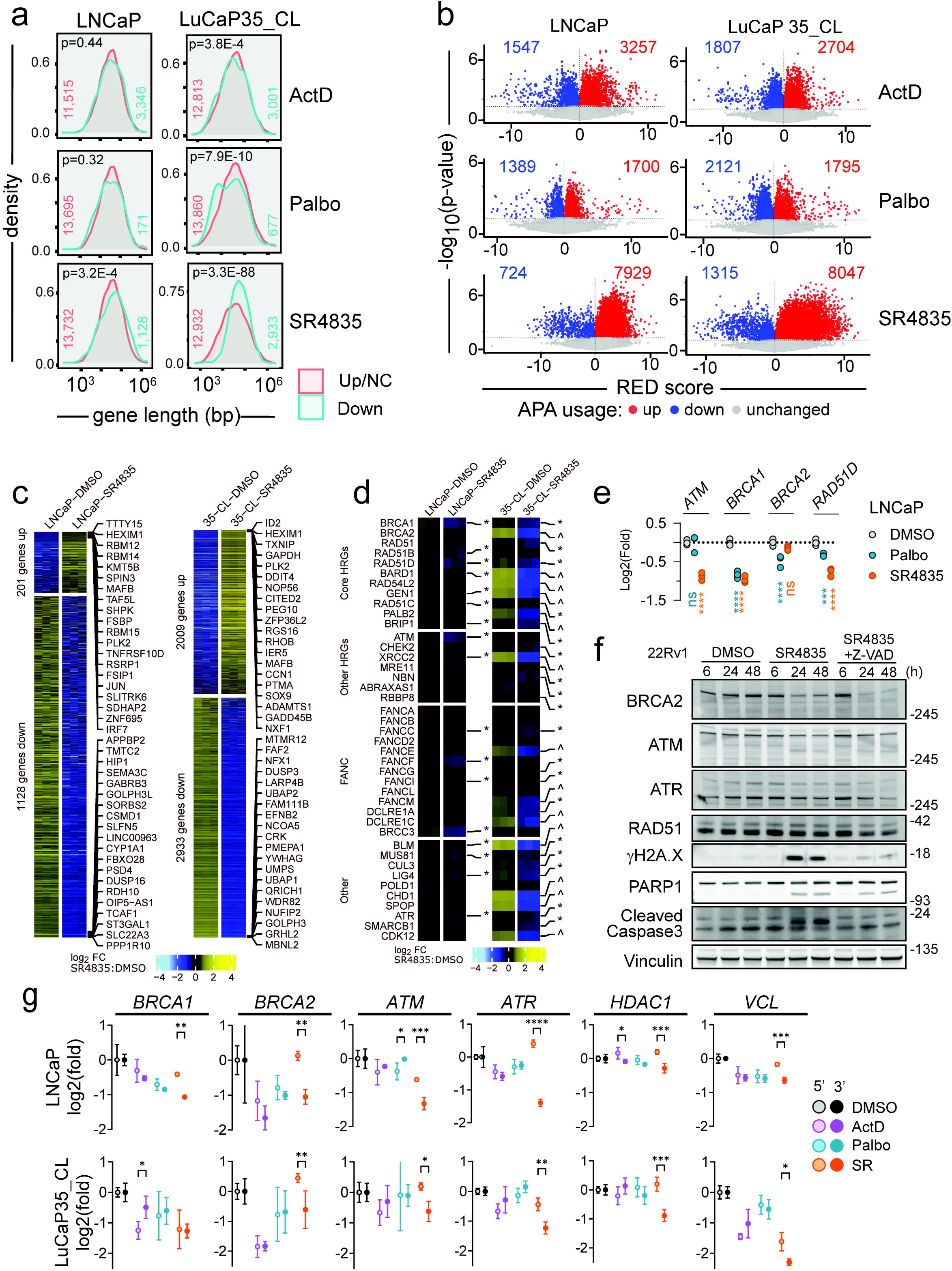
Acute CDK12 inhibition increases alternate polyadenylation usage and downregulation of DNA repair genes. **(a)** Genes downregulated by acute CDK12 inhibition skew longer. Distribution by gene length (genome span) (bp) of downregulated genes by RNA-seq (<-2 fold, FDR <0.05, ‘n’ depicted on plot) in LNCaP and LuCaP35_CL prostate cancer cell lines following six hours of exposure to vehicle (DMSO), CDK4/6 inhibitor palbociclib (Palbo, 10uM), broad RNA Pol-II inhibitor actinomycin D (ActD, 5ug/mL), or CDK12/13 inhibitor SR4835 (200nM) (n=3). Plots were made with ShinyGO 0.80 (86) and show significance (t-test) of downregulated vs upregulated (Up) or no change (NC) in expression genes. **(b)** Acute CDK12 inhibition increases intronic alternate polyadenylation site usage. APAlyzer analysis of the RNA-seq data showing treatment effect on intronic APA usage (compared to vehicle). Colored dots show significant (p<0.05) up and downregulated relative expression difference of APA sites. Numbers inside plots indicate the number of significant up (red) or down (blue) APA sites. **(c)** Treatment with the CDK12/13 inhibitor SR4835 alters gene expression. Heatmap showing log2(fold) color-coded expression and genes with >2 fold change up or down (FDR<0.05) upon SR4835 treatment with top and bottom 20 labeled. **(d)** Multiple HR genes are downregulated with SR4835 treatment. Heatmap as in (c) showing expression of HR-related genes upon acute CDK12 inhibition. The * indicates genes with any negative FC at FDR<0.05 and ^ indicates FC<-2 at FDR<0.05. **(e)** Some, but not all, HR gene downregulation with SR4825 is due to cell arrest. Example genes from the LNCaP RNA-seq showing the magnitude of Palbo vs SR4835 effects on RNA expression (log_2_(fold) RPKM inhibitor vs DMSO). Significance determined by two-way ANOVA with Dunnett multiple testing correction. **(f)** Acute CDK12 inhibition decreases BRCA2 protein and induces apoptosis after 24h. Western blot of 22Rv1 cells treated 6, 24, or 48 hours with vehicle (DMSO), SR4835 (200nM), or SR4835 plus pancaspase inhibitor Z-VAD (50µM). Lysates were probed for key DNA repair genes and markers of DNA damage (γH2A.X) and apoptosis (total/cleaved PARP1, cleaved caspase 3). Figure is a composite from two gels run with identical lysates (3-8% tris-acetate for BRCA2, ATM, ATR, Vinculin and 4-12% bistris for RAD51, γH2A.X, PARP1, and CC3). **(g)** Long genes show 5’/3’ transcript imbalance upon acute CDK12 inhibition. qPCR with same RNA samples as above (a-e) using two sets of primers for each target to show selective loss of 3’ transcript in long DNA repair-associated genes plus *Vinculin/VCL*, a long gene not involved in DNA repair. Plots show mean log2(fold) vs DMSO −/+ stdev (n=3) with significance between 5’/3’ primers determined by two-way ANOVA.

To determine if levels of proteins involved in DNA repair pathways were concordantly diminished, LNCaP and 22Rv1 prostate cancer cells were treated for 6, 24 or 48h with SR4835 and proteins were analyzed by immunoblot (**Fig 4f, Fig S4e,f**). In agreement with previous studies, BRCA2, ATM, and ATR protein decreased at 24h and 48h post treatment with 200nM SR4835. Acute CRISPR/Cas9 mediated CDK12 KO in LNCaP and C4-2B cells also led to decreased BRCA2 and ATM protein expression by immunoblot (**Fig. S4g**). Double strand DNA (dsDNA) breaks, as indicated by γH2A.X, increased by 48h (LNCaP) and 24h (22Rv1) of SR4835 treatment but was largely ablated with the addition of Z-VAD, a pan-caspase inhibitor. Similar results were observed with an ovarian cancer line, Skov3 (**Fig S4f**). Thus, while SR4835 does cause moderate decreases in key DNA repair proteins, most of the corresponding γH2A.X appears due to caspase-dependent apoptosis and not impaired DNA repair directly.

To test if transcript shortening was responsible for the DNA repair gene downregulation with SR4835, qRT-PCR was performed with specific primers for 5’ and 3’ regions of each target. SR4835, but not ActD or Palbo, led to specific 3’ transcript loss in the long genes tested including *BRCA1*, *BRCA2*, *ATM*, *ATR*, *HDAC1*, and *VCL* (**Fig 4g**). Together, these results confirm mRNA shortening, the APA activation phenotype, and downregulation of transcripts and proteins involved in HR in prostate cancer models under acute CDK12 loss conditions.

*CDK12* is classified as a ‘common essential’ gene (https://depmap.org/portal/) and CDK12/13 inhibitors cause apoptosis after 24-48h (**Fig 4f and S4e,f**). Despite this essentiality, a subset of human prostate cancers do tolerate the loss of *CDK12* (**Fig 1**). We next investigated the possibility that cells adapted to CDK12 loss might show a different phenotype than cells undergoing acute CDK12 depletion. To carry out these studies, we developed a new *in vitro* model of *de novo CDK12^BAL^* prostate cancer by generating a cell line (LuCaP189.4_CL) from the LuCaP189.4 patient derived xenograft (PDX) (51) that carries bi-allelic *CDK12* frameshift mutations (p.S345Gfs*10, p.S521Qfs*89) (**Fig S4h**). LuCaP 189.4 does not express detectable CDK12 protein (**Fig S4i**) and exhibits a classic tandem duplicator phenotype (TDP) that is a hallmark of *CDK12^BAL^* tumors (**Fig S4j**). We quantified the abundance of 5’ vs 3’ transcript levels by qRT-PCR in three *CDK12* intact PC models and LuCaP189.4_CL and found that only the *CDK12^BAL^* LuCaP189.4_CL displays a selective 3’ decrease (log2(fold) mean difference) in *ATM* (0.72, p<0.0001) and *ATR* (0.72, p=0.004), but non-significant or 3’ increases in *BRCA1*, *BRCA2, HDAC4,* and *VCL* (**Fig 5a**), indicating that these long genes are less affected by APA and splicing effects in cells naturally adapted to tolerate the absence of *CDK12*.

**Figure 5.**
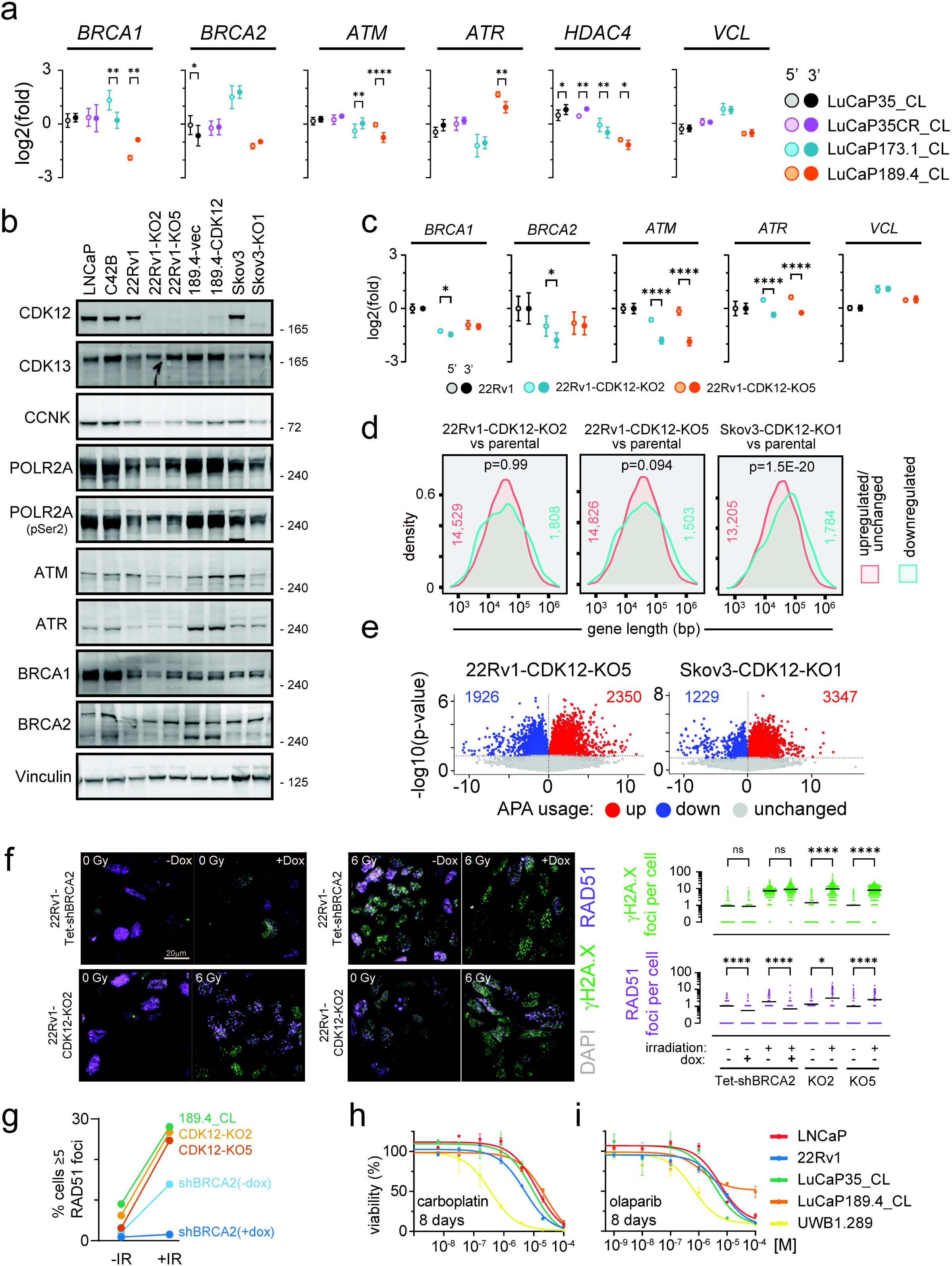
Cells adapted to CDK12 loss do not show dramatic HR gene downregulation and retain an intact HR pathway. **(a)** *CDK12^BAL^* cells have mostly restored 5’/3’ transcript balance. qPCR with CDK12-intact (Lu-CaP35_CL; LuCaP35CR_CL; LuCaP173.1_CL) and CDK12^BAL^ (LuCaP189.4_CL) prostate cancer cell lines using 5’ and 3’ targeting primers. Plot shows mean-centered log2(fold) −/+stdev (n=3), with significance determined by two-way ANOVA. **(b)** Cells with stable *CDK12^BAL^*show normal expression of key DNA repair genes. Immunoblot of factors involved in homology directed repair in prostate and ovarian cancer models with intact or absent CDK12. Figure is a composite from two gels run with identical lysates. **(c)** 22Rv1-CDK12-KO clones retain *ATM* 3’ loss but restore *BRCA1* and *BRCA2* transcript balance. qPCR as in (a) comparing 22Rv1 vs CDK12-KO clones. Mean log_2_(fold) −/+stdev (n=4) vs parental line with significance determined by two-way ANOVA. **(d)** 22Rv1 *CDK12* KO clones do not show selective long gene downregulation. Same analysis as in Fig. 4a. Plots show distribution by gene length of downregulated vs upregulated/unchanged genes in the indicated CRISPR KO clone line vs parental line. Significance was determined by t-test and ‘n’ is depicted on plots. **(e)** 22Rv1-CDK12-KO clones do not show increased APA usage. APAlyzer analysis (as in Fig. 4b) showing APA usage differences in 22v1-CDK12-KO5 and Skov3-CDK12-KO1 compared to parental cells. APA counts from 22Rv1-CDK12-KO2 and LuCaP189.4-CDK12 lines can be found in Fig. S6a. **(f)** *CDK12* loss does not impair RAD51 foci formation. 22Rv1 cells with Tet-inducible shBRCA2 were treated −/+ doxycycline (100ng/mL) for 4 days then exposed to 6Gy ionizing radiation (IR) and fixed at 3h post IR. The same was done with 22Rv1 CDK12 KO clones. Immunofluorescence staining was performed for γH2A.X and RAD51 and images were acquired by confocal microscopy. Left: representative images (white: DAPI, green: γH2A.X, purple: RAD51). Right: quantification of images (∼200-500 cells analyzed per treatment). Line is at mean and significance was determined by one-way ANOVA (Kruskal-Wallis). **(g)** Alternate quantification of (f) and Fig. S6c showing the percent of cells with 5 or more RAD51 foci before and after irradiation (IR). **(h-i)** *CDK12* loss does not confer platinum or PARPi sensitivity. Dose response curves for prostate cancer cell lines treated 8 days with carboplatin (h) or olaparib (i) (n=3, mean-/+stdev). UWB1.289 is a control *BRCA1*mut ovarian cancer line. Y-axis shows relative viability vs no drug and x-axis shows concentration (moles/L).

To further study the transcriptional features of cells adapted to *CDK12* loss, CRISPR-mediated knockout (KO) clones were generated in 22Rv1 (two clones: KO2 and KO5) and Skov3 cells (one clone: KO1), and LuCaP189.4_CL cells were engineered to re-express CDK12 (**Fig 5b**). Consistent with previous studies, very few clones tolerated *CDK12* deletion, and those that did grow slower than the parental lines (**Fig. S4k**). The 22Rv1 KO clones were generated with dual sgRNAs generating mutations in exons 1 and 4 that were validated by targeted PCR showing large insertions (**Fig. S5a**) and frameshift indels as seen from RNA-seq (**Fig. S5b**). The Skov3-CDK12-KO1 clone was generated with a single CDK12 sgRNA and contains two frameshift indels (+1 and −1) detected in exon 1 RNA-seq reads (not shown). At the protein level, cells with *CDK12* deletion showed slight increases in CDK13 and decreases in CCNK/CyclinK but no obvious decrease in p-Ser2 RNA Polymerase II/RBP1/POLR2A levels (**Fig 5b**). The *CDK12*-KO clones did not show decreases in ATR, BRCA1, or BRCA2 protein but did show decreased ATM compared to the parental lines (**Fig 5b**). Interestingly, the *CDK12^BAL^* LuCaP189.4_CL showed comparable levels of these DNA repair genes, and although the re-expression of CDK12 was not very robust, ATM protein was slightly increased (**Fig 5b**). Consistent with the results in the *de novo CDK12^BAL^* LuCaP189.4_CL, the 22Rv1 *CDK12*-KO clones also exhibited persistent 3’ vs 5’ transcript decreases (mean log2(fold)) in *ATM* (1.16, p <0.001 for KO2 and 1.71, p<0.0001 for KO5) and *ATR* (0.84, p<0.0001 for KO2 and 0.88 p<0.0001 for KO5), but minimal 5’/3’ difference in *BRCA1*, *BRCA2,* or *VCL* (**Fig 5c**). The 22Rv1 *CDK12*-KO clones did show lower overall *BRCA1* (–1.46 KO2, –1.01 KO5) and *BRCA2* (–1.79 KO2, –0.97 KO5) mRNA levels (log2(fold) vs DMSO, 3’ primer set) (**Fig 5c**). However, reduced transcripts of these genes may be due to cell-cycle linked expression and the slower proliferation of these clones (**Fig S4k**) and not a direct result of *CDK12* loss. The 22Rv1-CDK12 KO clones did not display an obvious TDP pattern from low coverage WGS copy number analysis (**Fig. S5c**), especially compared to LuCaP189.4 (**Fig. S4j**).

RNAseq based assessments of the isogenic *CDK12* intact versus knockout lines identified no enrichment for longer genes among those with lower expression in the stable 22Rv1-*CDK12*-KO clones (p=0.99 for KO2 and p=0.094 for KO5), though there was enrichment in the Skov3-*CDK12*-KO1 line (p=1.5e^-20^) (**Fig 5d**). APA analysis found modest increases in IPA site usage (UP/DOWN ratio) in the *CDK12*-KO5 (1.2) and Skov3-*CDK12*-KO1 (2.7), but not 22Rv1-*CDK12*-KO2 (0.8) (**Fig 5e and S6a**), far less dramatic than under acute CDK12 inhibition which had UP/DOWN ratios of 11.0 in LNCaP and 6.1 in LuCaP35_CL (**Fig 4b, S4b**). These results show that, with the notable exception of *ATM*, most long genes (including *BRCA1* and *BRCA2*) do not show downregulation in tumor cells that have adapted to *CDK12* loss. Though some genes show IPA alterations, the phenotype is far less apparent than under acute loss conditions. Furthermore, though CDK12 inactivation in the Skov3 ovarian cancer cells showed some preferential downregulation of long genes overall (**Fig 5d**), BRCA1 and BRCA2 protein were not affected (**Fig 5b).**

### Cells adapted to *CDK12* loss are HR competent and lack exceptional sensitivity to PARPi or platinum chemotherapy

A key early step in HR involves BRCA2-mediated loading of RAD51 onto resected ssDNA. Loss of key HR pathway members, including BRCA1, BRCA2, or PALB2, all lead to loss of RAD51 loading and initiation of HR repair (52). Though CDK12-KO cells retain BRCA1 and BRCA2 protein expression (**Fig 5b**), it remains possible that HR function could still be altered by other means. To test this possibility, in addition to the CDK12-KO clones, LNCaP, 22Rv1, and Skov3 cells were engineered with Tet-inducible shBRCA2 or shCDK12 (**Fig S6b**). Cells were exposed to 6Gy ionizing radiation (IR) and immunostained for γH2A.X and RAD51 at 3h post irradiation (**Fig 5f and S6c-e**). 22Rv1-Tet-shBRCA2 cells functioned as an HRd control: shBRCA2 without dox went from 1.0 to 1.8 RAD51 foci per cell after IR, while dox treated cells went from 0.5 to 0.6 (**Fig 5f**). Following IR, the RAD51 foci per cell in 22Rv1-CDK12-KO lines increased from 1.2 to 2.9 in KO2 (p=0.04) and 0.9 to 2.4 in KO5 (p<0.0001) (**Fig. 5f**). Likewise, the *CDK12^BAL^* LuCaP189.4_CL was competent at inducing RAD51 foci (1.4 to 5.4 foci per cell, p<0.0001) (**Fig S6c**). Since only G2/M cells can use HR repair, we also used an alternate quantification metric of % cells with 5 or more foci (i.e. % of population undergoing HR) which showed that *BRCA2* knockdown greatly reduced the RAD51 population after IR (13.9% indox, 1.5% in +dox) while the 22Rv1-*CDK12*-KO clones and LuCaP189.4_CL showed a high proportion (25-28%) of HR-functioning cells (**Fig 5g**). Additional experiments with Tet-shBRCA2 and Tet-shCDK12 in LNCaP (**Fig S6d)** and Skov3 (**Fig S6e**) produced similar results, with CDK12 knockdown unable to prevent RAD51 foci induction (i.e. non-significant differences after IR between shCDK12 line −/+dox). These results with CDK12 knockdown and knockout, across multiple models, show that CDK12 deficient cells maintain the ability to load RAD51 and initiate HR repair.

Though cells adapted to survive CDK12 loss do not exhibit compromised HR, the possibility remains that CDK12 loss could sensitize to platinum chemotherapies or PARPi via other mechanisms (53–55). Dose response curves were performed with carboplatin (**Fig. 5h**) and the PARPi olaparib (**Fig. 5i)** using various prostate cancer lines, plus *BRCA1*-deficient UWB1.289 ovarian cancer cells as a *bona fide* HRd control (56). Though UWB1.289 showed the expected sensitivity to both carboplatin (EC50 0.36uM) and olaparib (0.71 uM) at 8 days, LuCaP189.4_CL was not sensitive (carboplatin EC50 = 19.15; olaparib EC50 >100uM). Inducible knockdown of *BRCA2* in 22Rv1 cells altered the EC50 of carboplatin from 5.92 μM to 2.16 μM with dox, whereas the two 22Rv1 *CDK12*-KO clones showed either a greater or equivalent EC50 (**Fig S7a)**. Skov3-*CDK12*-KO1 showed no difference in response to carboplatin (**Fig. S7b**) or olaparib (**Fig. S7c**) at 8 days treatment compared to control Skov3 cells. The 22Rv1-*CDK12*-KO clones and *CDK12^BAL^* LuCaP189.4_CL showed mixed responses to PARPi: in a 12-day treatment, LuCaP189.4_CL (EC50 0.20μM) and the 22Rv1-*CDK12*-KO clones (0.88μM for KO2, 0.92μM for KO5) displayed some sensitivity to olaparib compared to 22Rv1 (17.37μM), but not to the same extent as UWB1.289 (0.08μM) (**Fig S7d).** LuCaP189.4_CL did not show enhanced sensitivity in an 8 day exposure to rucaparib (**Fig S7e**). In 14 day treatments with organoids harvested from PDX tumors, *CDK12*-intact LuCaP lines (23.1, 170.2, 86.2CR) had EC50 ranges of 2.23-6.33μM for olaparib while LuCaP189.4 was at 13.30μM, showing no apparent sensitization (**Fig S7f**) as was also the case with rucaparib (**Fig S7g**). Collectively, these results indicate that stable CDK12 loss does not sensitize to carboplatin, though the effect on PARPi sensitivity is less consistent with clonal variation, but significantly less than the *bona fide* HRd line UWB1.289.

### CDK13 is synthetic lethal in cells with biallelic CDK12 loss

CDK12 and CDK13 have overlapping and unique roles in regulating transcription and RNA metabolism, and both function in a heterodimer with Cyclin K (CCNK) (57). Analysis of CRISPR screen data from the Dependency Map project (https://depmap.org/portal/) shows that CDK13 depletion is generally tolerated in most lines, while CDK12 loss is detrimental in most cells (**Fig 6a**). Moreover, CCNK depletion has an even more negative fitness effect (<-1 CERES score). The LuCaP189.4_CL, which does not express CDK12, was the only natural line tested without negative growth effects upon sgCDK12 transduction (**Fig. 6b**), while LNCaP (**Fig. 6c-d)**, C4-2B (**Fig. 6e**), Skov3 (**Fig. 6f**), and 22Rv1 (**Fig. S7h,i**) all showed significantly reduced growth (p<0.006) by confluence in sgCDK12 vs sgAAVS1, most dramatically in C4-2B (47.3% with sgAAVS1, 2.6% with sgCDK12). While LNCaP, C4-2B, and Skov3 tolerated CDK13 CRISPR lentivirus with no significant difference vs AAVS1 (**Fig 6c-f**), cells with stable *CDK12^BAL^* loss: LuCaP189.4_CL and Skov3-CDK12-KO1, showed a marked growth inhibition with sgCDK13 vs control (LuCaP189.4_CL: 59.5% vs 88.3%, p<0.0001, Skov3-KO1: 5.6% vs 12.9%, p=0.004) (**Fig 6b,g**). Of interest, the growth of 22Rv1 was repressed by either CDK12 or CDK13 sgRNAs (**Fig S7i**). However, 22Rv1-*CDK12*-KO5 appeared to have almost complete growth suppression with sgCDK13 from Day 5 to 14 (0.8% to 0.8%) compared to sgAAVS1 (0.6% to 5.5%) while parental 22Rv1 with sgCDK13 still had measurable growth (1.6% to 6.3%) (**Fig S7h-j**).

**Figure 6.**
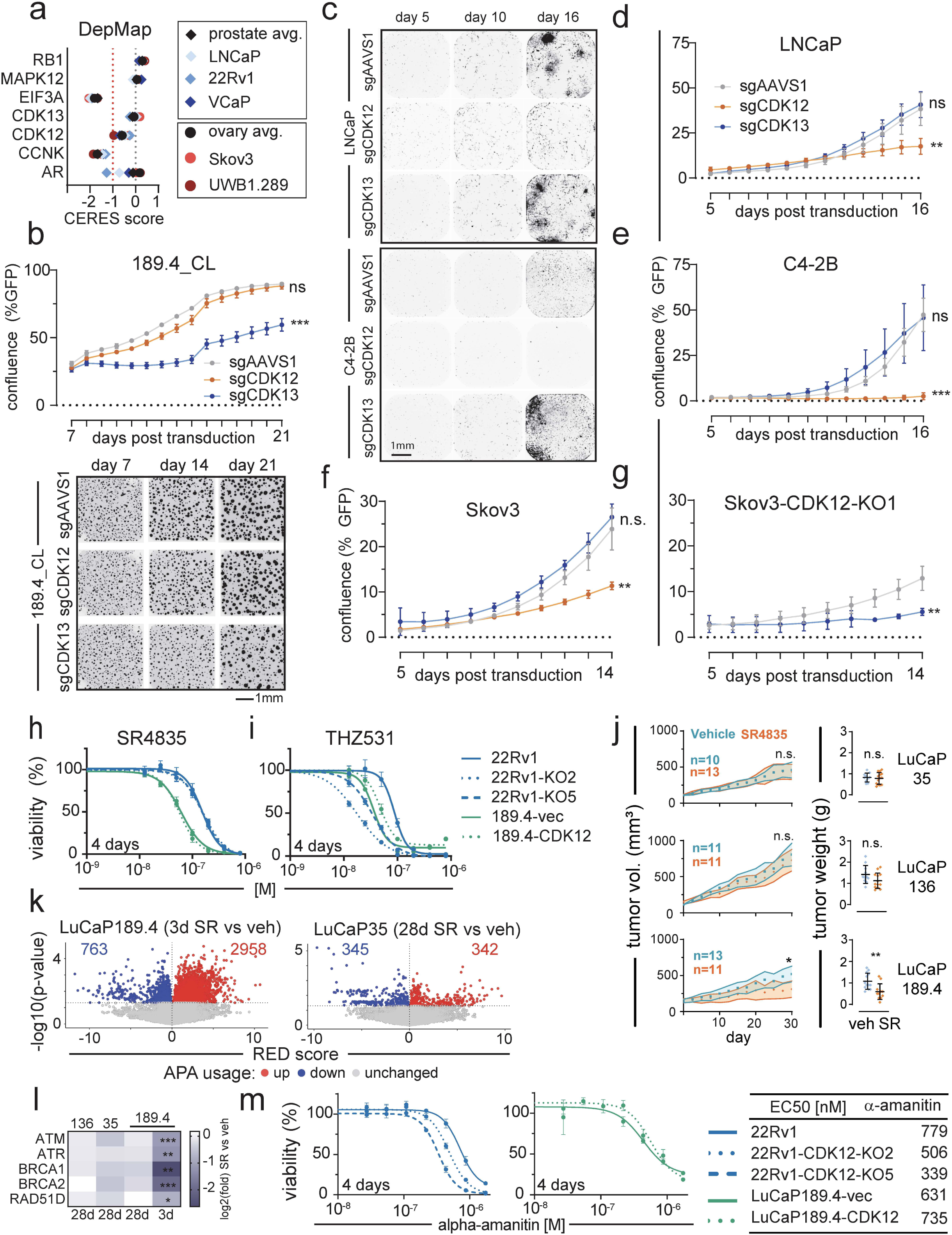
Prostate cancers with CDK12^BAL^ are sensitive to CDK13 loss and therapeutics targeting transcription. **(a)** CDK12 loss is generally detrimental to cells. Selected control and CDK12-related sgRNA fitness results from DepMap for relevant cell lines and average score across cancer type. RB1 is a control for positive selection, MAPK12 (p38γ) is a control for neutral selection, and EIF3A is a control for negative selection. **(b)** LuCaP189.4_CL growth is greatly impaired by sgCDK13. GFP-tagged LuCaP 189.4_CL were transduced with CRISPR/Cas9 vectors containing dual sgRNAs against PPP1R12C/AAVS1 (neg. control), *CDK12*, or *CDK13* and growth was monitored by GFP imaging starting on day 7 (n=5). Plot shows mean confluence (%GFP −/+stdev, n=5) with significance vs sgAAVS1 determined by two-way ANOVA. Images below graph show example microscopy from days 7, 14, and 21. **(c-e)** *CDK12* KO is poorly tolerated in LNCaP and C4-2B cells. GFP-tagged LNCaP and C4-2B were transduced with dual sgRNA vectors and monitored as in (b). **(c)** Example images showing sgRNA effect on cell growth and growth rate plots (mean confluence (%GFP) −/+stdev, n=5) over time for LNCaP **(d)** and C4-2B **(e)** with significance vs sgAAVS1 determined by two-way ANOVA. **(f-g)** *CDK13* sgRNA is detrimental in Skov3 lacking *CDK12*. Skov3 **(f)** or Skov3-*CDK12*-KO1 **(g)** were transduced with sgRNAs and monitored as in (c-e). Plot shows mean confluence (%GFP) −/+stdev, n=5) with significance vs sgAAVS1 determined by two-way ANOVA. **(h-i)** *CDK12* loss corresponds with sensitivity to CDK13 inhibition. Plots show dose response curves for 22Rv1 or 22Rv1-CDK12-KO clones and 189.4 empty vector (189.4-vec) or CDK12-transduced (189.4-CDK12) treated four days with SR4835 **(h)** or THZ531 **(i)** CDK12/13 inhibitors. Y-axis shows relative viability vs no drug and x-axis shows concentration (moles/L). (**j**) *In vivo* LuCaP189.4 tumors respond to SR4835. PDX LuCaP lines were treated 28 days with vehicle or SR4835 (20mg/kg, 5 days on, 2 days off). Plots show number of mice per group, tumor volume (mean with 95%CI) over time, and final tumor weight. Significance was determined by Kolmogorov-Smirnov test (tumor volume) or unpaired, two-tailed t-test (tumor weight). (**k**) Three day SR4835 treatment *in vivo* increases APA usage, but the effect is absent in 28 day tumors. APAlyzer analysis (as in Fig. 4b) on RNA-seq from the treated PDX tumors harvested in (j) and LuCaP189.4 treated 72h (n=3 tumors per treatment). (l) Heatmap showing mean log_2_(fold) expression of genes involved in HR determined by RNAseq from tumors resected after 3d or 28d treatment with SR4835 compared to vehicle, as in (k). (**m**) CDK12 loss increases sensitivity to α-amanitin. 22Rv1 and LuCaP189.4_CL were treated four days with α-amanitin to generate dose response curves, as in (h,i). Plots shows mean-/+stdev(n=4) and table shows effective concentration 50% viability (EC50) values and legend.

Due to high protein conservation, all currently available pharmacological inhibitors of CDK12 also inhibit CDK13. We performed dose response curves with two different selective CDK12/13 inhibitors: SR4835, an ATP competitive inhibitor (21) (**Fig 6h and S7k**), and THZ531, a covalent binding inhibitor (28) (**Fig 6i and S7l-m**). SR4835 was reported to have a dissociation constant of 4.9nM for CDK13, 98nM for CDK12, and >800nM for all other kinases tested (21) while THZ531 was measured to have an IC50 value of 69nM for CDK13, 158nM for CDK12, and >8µM for CDK7 and CDK9 (28). LuCaP189.4_CL and the *CDK12*-KO 22Rv1 clones all showed higher sensitivity to THZ531 with EC50 of 88.6nM for 22Rv1 vs 17.3nM (KO2) and 33.3nM (KO5) and 37.8nM (LuCaP189.4-vec) vs 52.6nM for cells with re-expressed CDK12 (LuCaP189.4-CDK12) (**Fig 6i)**. Under 3D growth conditions, LuCaP189.4 spheroids were also more sensitive to THZ531 (EC50: 77nM) than LuCaP35 (145nM) or RWPE1 (223nM) (**Fig. S7m-n**). Surprisingly, the *CDK12*-KO clones did not show the same curve shift with SR4835 (**Fig 6h)**, though LuCaP189.4_CL was quite sensitive with an EC50 of 38nM (**Fig 6h)** and 29nM (**Fig S7k,n**) measured in different experiments. This difference could be due to the fact that SR4835 is a non-covalent inhibitor, while THZ531 is a covalent modifier. Unfortunately, THZ531 has not been deemed usable for *in vivo* use while SR4835 has been used previously in mice (21).

To confirm whether the CDK13 vulnerability could exploited for *in vivo* treatment, we performed xenograft drug treatments using three LuCaP PDX lines (LuCaP35, LuCaP136, and LuCaP189.4) treated for 28 days with vehicle or SR4835. The dose schedule was the same as previously established in a report from the developers of the compound in a breast cancer PDX experiment (21). LuCaP 35 and 136 are both *CDK12*-intact. At the final 28 day timepoint, vehicle treated LuCaP 189.4 tumor volumes (mm^3^[95%CI]) measured 521[411-630] while SR4835 tumors were smaller at 310[196-425] with a significant (p=0.046) cumulative difference between growth curves (Kolmogorov-Smirnov, twotailed) (**Fig 6j**). Tumors were harvested and weighed, with SR4835 treated tumor average weight being significantly smaller (0.59 vs 1.08 g, p=0.0015, Mann-Whitney test) (**Fig 6j**). There were no significant tumor volume or weight reductions in the *CDK12*-intact lines. Taken together, these results support the hypothesis that cells lacking CDK12 become dependent on CDK13 for their CCNK/CyclinK activity, thus presenting a potential targeted vulnerability with potential *in vivo* efficacy, even with dual CDK12/13 targeting compounds.

We performed RNA-seq on three of the harvested tumors per group after 28 days SR4835, plus a set of LuCaP189.4 PDX tumors treated acutely for 3 days with SR4835 or vehicle. Results were analyzed by APAlyzer and found large upregulation of APA sites in LuCaP189.4 after 3 day treatment (2,958 Up) but not LuCaP 35 after 28 days treatment (342 Up) (**Fig. 6k**). Furthermore, the UP/DOWN ratio of APA usage was only >1.0 for LuCaP189.4 at the 3 day treatment point (3.9 ratio) and at or below 1.0 for all 3 lines in the 28 day treated tumors (**Fig. S7o**). A selection of example long DNA repair gene transcripts were significantly decreased with the 3d treatment including: *ATM* (log_2_ FC –1.1), *ATR* (−1.2), *BRCA1* (−2.6), *BRCA2* (−2.0), and *RAD51D* (−0.8)(all p<0.01 in SR4835 vs vehicle) (**Fig 6l**). None of these genes were significantly decreased in the 28 day treated tumors, suggesting that cells able to survive and grow in the presence of the CDK12/13 inhibitor no longer show the long gene APA phenotype and downregulation.

### CDK12 loss increases sensitivity to selected therapeutics targeting transcription

The most studied function of CDK12 is to maintain RNA polymerase II processivity and proper splicing and polyadenylation. Tumors adapted to CDK12 loss show modest if any alteration in the transcript lengths or APA usage of large genes involved in HR (**Fig. 3g and 5b-d**) and no substantial change in RNA-Pol II Ser2 phosphorylation (**Fig. 5b**), but these adapted cells may still have impaired transcriptional processes which could be exploited. We next tested if cells with CDK12 loss may exhibit enhanced sensitivity to therapeutics targeting transcription mechanisms. Dose response curves revealed a modest selective sensitivity in *CDK12^BAL^* lines to α-amanitin, an RNA-Pol II poison with EC50 of 779nM for 22Rv1 compared to 506nM (KO2) and 339nM (KO5) (**Fig 6m**). Re-expression of CDK12 in the LuCaP189.4_CL promoted amanitin resistance to a small but not statistically significant degree (631nM to 735nM, p=0.4) (**Fig 6m**). These results suggest that cells adapted to CDK12 loss may continue to exhibit compromised transcriptional mechanisms, and that this may lead to a vulnerability to amanitin-class agents or other drugs that target mRNA synthesis or processing.

### CDK12 loss does not consistently increase sensitivity to WEE1, ATR, or CHEK1 inhibitors

Among other potential targetable DNA repair pathway vulnerabilities considered, we tested two WEE1 inhibitors: (adavosertib/MK-775) (**Fig S8a**) and PD0166285 (**Fig S8b**). The EC50 values for LuCaP189.4_CL (with or without CDK12 re-expressed) and 22Rv1-*CDK12*-KO2 were all greater than parental 22Rv1 (**Fig S8c**). However, 22Rv1-*CDK12*-KO5 did show some increase in sensitivity (EC50) compared to the parental line to adavosertib (0.04μM vs 0.11μM) and PD0166285 (0.10μM vs 0.37μM) (**Fig S8c**). Considering that adapted *CDK12^BAL^* cells do retain some ATM transcript shortening and protein decreases (**Fig 5a,b**), we evaluated ATR as a potential vulnerability. We tested two ATR inhibitors, berzosertib (**Fig S8d**) and elimusertib (**Fig S8e**), but found that LuCaP189.4_CL had the highest EC50 values to both inhibitors (**Fig S8f**). 22Rv1-*CDK12*-KO5 (but not KO2) showed a slight increase in sensitivity to berzosertib, and although both KO clones showed some increased sensitivity to elimusertib vs parental, the EC50 curves were similar to LNCaP and C4-2B reducing the likelihood of a CDK12-specific effect (**Fig S8f**). Lastly, we tested two CHEK1 inhibitors, rabusertib (**Fig S8g**) and MK-8776 (**Fig S8h**), as potential ATR-related dependencies. However, the results show that LuCaP189.4_CL was the least sensitive line tested to both (**Fig S8i**), and only 22Rv1-*CDK12*-KO5 showed a clear increase in sensitivity vs parental to both rabusertib (0.53μM vs 1.81μM) and MK08776 (0.48μM vs 1.54μM). In summary, one of the tested *CDK12* loss models (22Rv1-*CDK12*-KO5) showed modest CDK12-associated growth repression by WEE1, ATR, and CHEK1 inhibitors. However, since differential treatment effects were not shared with 22Rv1-*CDK12*-KO2 or Lu-CaP189.4_CL it appears that these sensitivities are not consistent, but may relate to alternative mechanisms by which cells survive CDK12 loss.

## DISCUSSION

In the present work we sought to identify a molecular basis for the discrepancy between the poor clinical responses to PARPi in patients with *CDK12* alterations and preclinical studies demonstrating that *CDK12* loss or pharmacological inhibition compromises HR, phenocopies ‘BRCAness’ and results in synthetic lethality to drugs impeding PARP function. A central feature of most prior studies evaluating CDK12 and HR is the conduct of very short term *in vitro* experiments, with timepoints usually less than 72 hours after pharmacological inhibition of CDK12 activity or repressing CDK12 by genetic methods (12, 16, 19, 21, 26, 28, 58). A key reason for evaluating such acute timepoints is largely due to the fact that CDK12 inhibition leads to proliferation arrest and/or cell death, as CDK12 is a common essential gene in most cells (15, 16, 28, 59). Acute loss of CDK12 activity clearly results in increased APA site usage and the diminished expression of a cohort of large genes, including several involved in HR. This consequence can contribute to HRd in the immediate setting, and act in synergy with PARPi and DNA damaging agents (26, 29, 60–62). However, it should be noted that cell cycle arrest also results in the downregulation of many HR-associated genes, and it is not straightforward to attribute causality specifically to CDK12-mediated transcriptional compromise versus cell cycle-regulated expression (50). We note that the CDK4/6 inhibitor palbociclib caused more significant downregulation of DNA repair pathways than the CDK12/13 inhibitor SR4835.

In contrast to the acute effects of functional CDK12 loss, cells naturally adapted to survive and proliferate in the context of *CDK12* absence, such as the LuCaP189.4 model and human tumors, show less differential down-regulation of long genes compared to acute CDK12 repression. In analyses of human tumors with *CDK12^BAL^*, the expression of genes regulating HR are not reduced, and these tumors do not exhibit genomic scars (such as high LOH) or mutation signatures (such as COSMIC mutation signature 3) that reflect compromised HR found in tumors with *BRCA1* or *BRCA2* loss. Further, using functional assays, such as RAD51 foci formation after radiation, both *de novo CDK12^BAL^* cells, and those adapted to tolerate CDK12 deletion by CRISPR/Cas9, demonstrate HR competency.

Through literature searches and cell line database queries, we were not able to identify any cancer models with *de novo CDK12^BAL^*, and only one prior study where a stable *CDK12* deletion in a cancer cell line was generated (63). This supports the conclusion that *CDK12* is a common essential gene in most cells, with a severe impact on viability that is challenging to overcome. In agreement with the results presented here, the above mentioned A2780 ovarian cancer cell line model with stable *CDK12* deletion showed markedly slower proliferation, increased rates of apoptosis, and no significant reductions in HR-related proteins including ATM, BRCA1, ATR or FANCD2. The A2780 *CDK12* knockout line was also not sensitive to platinum chemotherapy or olaparib, and was 5-fold more resistant to the ATR inhibitor VE822 and the WEE1 inhibitor MK1775 (63). We observed variable sensitivities to WEE1, ATR, and CHEK1 inhibitors in the context of CDK12 loss, suggesting actionable vulnerabilities in specific *CDK12* loss-adapted contexts which have yet to be characterized.

Though we did see modest sensitivity to PARPi in *CDK12^BAL^*cells in a few experiments, the lack of consistent HR gene downregulation, no effect on RAD51 foci formation, and low scoring of HRd genomic signatures, indicate that PARPi as a monotherapy is unlikely to benefit patients with CDK12^BAL^ tumors by exploiting HRd. However, other aspects of compromised CDK12 function could provide a therapeutic window for PARP inhibition as well as other targets. Recent studies revealed that PARPi can act beyond HR and affect other critical cell functions where CDK12 activity may be important including transcription, RNA splicing, R-loop repair, and replication (48, 53, 55, 64–67).

The identification of pharmacological vulnerabilities conferred by CDK12 loss is an important research priority. While we did not observe consistent effects using ATR or WEE1 antagonists, CDK13 appears to be an ideal synthetic lethal target for *CDK12^BAL^* cells, as complete loss of cyclin K activity in tumors would be severe, and other tissues could likely tolerate CDK13 inhibition in the setting of functional CDK12 (12). However, available pharmacologic inhibitors have been unable to separate CDK12 and CDK13 antagonism due to high protein similarity. This can lead to some toxicity concerns, especially with covalent inhibitors (68). However, CDK12/13 dual inhibitors may still be useful due to the enhanced sensitivity of *CDK12^BAL^* tumors to CDK13 inhibition. Furthermore, though less effective than THZ531 in culture assays, SR4835 was effective and generally tolerated well in mice and has been used in at least two published studies (21, 69). We observed modest differential sensitivity in several *CDK12^BAL^* models to the transcriptional inhibitor α-amanitin. Amanitin-based compounds are extremely toxic, but there is interest in using these drugs in antibody drug conjugates (ADCs)(70), which would offer a way to deliver toxic payloads directly and selectively to the tumor.

In addition to transcription-related consequences of CDK12 loss, it is very possible that CDK12 could play additional roles beyond RNAP II that have yet to be explored. Known direct substrates of CDK12 are fairly limited, with RNAP II being the most well studied. However, there is a report that 4EBP1 is a direct CDK12 substrate and confers a role in translation (71), and LEO1, part of the transcription elongation complex, was also reported as a direct CDK12 substrate (20). CDK12 has also been pulled down in complex with various other factors, including splicing regulators, though it is unclear if they are direct substrates (13, 15).

Of importance, the direct oncogenic role(s) of *CDK12* loss remains to be determined. Of interest is the limited organ site distribution of tumors with *CDK12* alterations. While the genomic structures of TDs produce a substantial increase in gene rearrangements, we found that very few of these are recurrent, and thus unlikely to serve as drivers of neoplasia. In contrast, a large number of oncogenes are contained within regions of tandem duplication of which several were recurrently gained in copy number, and could individually or collectively represent pathways driving cancer development. Notable among these recurrent events is the AR, which is either gained or amplified within a TD in 46% percent of *CDK12^BAL^* tumors. Of interest is the finding that no mCRPCs with a neuroendocrine phenotype had *CDK12^BAL^*, suggesting that *CDK12* loss is incompatible with neuroendocrine differentiation, or the *AR* locus copy gains serve to maintain AR signaling and prevent transdifferentiation to other lineages.

## METHODS

### Somatic mutation analysis

For whole exome sequencing (WES) data, single nucleotide variant (SNV) calling was performed using MuTect2 (GATK version 4.1.8.1), Strelka2 version 2.9.2 and VarScan2 version 2.4.4. Insertion and deletion (Indel) calling was performed using SvABA (commit 9813e84, https://github.com/walaj/svaba). All SNV and Indel calls were annotated using ANNOVAR (release 20200607). SNVs were included if they were detected by two or more callers, were not labeled by ClinVar significance as ‘benign’ or ‘likely_benign’, and were annotated as either ‘exonic’ or ‘splicing.’ Indel calls were included if they were not labeled by CLNSIG as ‘benign’ or ‘likely_benign’, were annotated as either ‘exonic’ or ‘splicing,’ and passed the following filters: Alt_reads_in_Tumor >= 5, Ref_reads_in_Tumor >= 10, and a variant allele frequency (VAF) of > 0.1. For patients with discrepant SNV or Indel calls between samples, manual curation was performed using IGV (version 2.16.2). For SU2C-WC WGS data, we obtained mutation calls from the original published results by Quigley et al. 2018 (3). For the Hartwig Medical Foundation (HMF) WGS data, the mutation callset was obtained via controlled access as part of Data Access Request DR-250. The liftOver tool was used to convert mutations calls from GRCh37 to GRCh38 coordinates, and these results were annotated with ANNOVAR (release 20200607), and mutations were included if they were not labeled by ClinVar significance as ‘benign’ or ‘likely_benign’, and were annotated as either ‘exonic’ or ‘splicing.’

### Germline mutation analysis

Germline mutation calling was performed for UW and SU2C-WC cohorts using GATK’s HaplotypeCaller using default settings. For WES data, the probe-set specific bait file was provided per sample as well. Output SNVs were annotated using ANNOVAR (release 20200607) and were included if they were annotated by CLNSIG as ‘pathogenic’ or ‘likely_pathogenic’. Manual curation was performed using IGV (version 2.16.2). Only somatic calls were provided in the provided data for HMF.

### Somatic copy number alteration analysis

For WES samples (SU2C-I and UW cohorts), the standard tumor-normal paired workflow of TITAN (v1.15.0) for WES was used [https://github.com/gavinha/TitanCNA]. Briefly, read counts were computed at 50 kb bins overlapping the exome bait set intervals.

Centromeres were filtered based on chromosome gap coordinates obtained from UCSC for hg38. The read coverage in each bin across the genome was corrected for GC content and mappability biases independently for tumor and normal samples using ichorCNA v0.3.4. Heterozygous SNPs were identified from the matched germline normal sample using Samtools mpileup. Only SNPs overlapping HapMap3.3 (hg38) were retained. The reference and non-reference allele read counts at each heterozygous SNP were extracted from the tumor sample. SNPs were not analyzed in chromosome X. Copy number analysis was performed using TitanCNA R package v1.15.0. Automatically generated optimal solutions were used.

The following methods were used for analyses of whole genome sequencing data. For SU2C-WC WGS data, copy number alterations, tumor purity, and tumor ploidy was predicted using TITAN (v1.15.0) as originally reported in Zhou et al. 2022 (72) using the workflow provided in https://github.com/GavinHaLab/TitanCNA_SV_WGS. For HMF WGS data, the copy number results were obtained via controlled access as part of Data Access Request DR-250. The results were originally analyzed using GRCh37 (hg19) reference genome.

Tumor cellularity was estimated by TITAN for each sample, and only samples with ≥ 20% tumor cellularity were included in downstream analyses.

### Mutational signature analysis

Mutational signature proportions were determined using the Analyze.cosmic_fit function from SigProfilerAssignment (v0.0.33) and Cosmic Version 3.4. Only SNV calls which passed our filtration were used for signature analysis. One input VCF was made per sample, with each input being reformatted to have five columns in the following order: chromosome, genomic coordinate, sample ID, reference base call, and alternative base call. When running Analyze.cosmic_file, the ‘build’ option was set to ‘GRCh38’, ‘input_type’ was set to ‘vcf’, and the ‘exome’ option was set to true for WES samples. All other options were left as default. To best capture HRD-related mutational signatures, we combined Cosmic Signatures SBS3 and SBS8 into an HRD-signature. Cosmic Signature 3 proportion was obtained by dividing the number of SBS3 SNVs by the total number of input SNVs per sample. The reported proportions were obtained by summing the counts of SBS3 and SBS8 per sample, then dividing by the total number of calls for each sample. We only considered SBS3 or SBS8 calls for samples with more than 50 passing SNVs and a combined SBS3 and SBS8 proportion of >0.05.

### Structural variation analysis

For samples based on short-read WGS, SvABA was used in tumor-normal paired mode for SV detection with default parameters. Intra-chromosomal SV events with span > 1 kb were retained. The SvABA workflow can be accessed at https://github.com/GavinHaLab/Ti-tanCNA_SV_WGS

### Classification of structural variants in mCRPC

SV types were annotated based on orientations of breakpoints and bin-level copy-number around breakpoints, as described previously (72). The copynumber near each breakpoint was evaluated using 10 kb bins. For each SV event, copy-number values of the bins located to the upstream and downstream of breakpoint 1 were denoted as *c_1_^up^* and *c_1_^down^*, respectively; similarly, the copy-number values for breakpoint 2 were denoted as *c_2_^up^* and *c_2_^down^*. In addition, then mean copy-number *c^mean^* of the 10 kb bins between the two breakpoints of the SV event were considered during SV classification. Intra-chromosomal SV events, *i.e.*, both breakpoints were located on the same chromosome, were classified into the SV types: deletion, tandem duplication, inversion, balanced rearrangement (intra-chromosomal), unbalanced rearrangement (intra-chromosomal). Interchromosomal SV events are classified as translocations.

### Tandem duplicator phenotype classification

Simple tandem duplications (TDs) were predicted from the copy number results genome-wide for each sample. For whole exome sequencing, duplication events must meet these criteria: ***(i)*** segment is shorter than 10 Mb; ***(ii)*** have flanking segments with lower copy number; (iii) the difference in copy number between the left and right flanking segment is ≤ 1. Out of the 632 WES samples, 581 were successfully analyzed by TITAN and having tumor purity >0.2 and *MAD* <0.25 were used to identify duplications. For WGS data, the tandem duplications were taken from the intersection of TITAN copy number segments and tandem duplication breakpoints from the final SV call set. As described previously (73), we used the Nearest-Neighbor Index (NNI) metric to distinguish the pattern of tandem duplications being dispersed (value near 1) as opposed to clustered. We applied a threshold for the NNI score as a guideline for manual curation of individual samples. For WES samples, TDP+ was defined as having NNI >1 and median segment length >100kb.

For WGS samples, TDP+ was defined as having NNI >1.25 and median segment length >100kb. Further manual inspection of the copy number profiles confirmed TDP+ and TDP-status.

### Tandem duplication size group analysis

To categorize TDP+ cases into TD size (i.e. length) groups, we applied Gaussian mixture model fitting using the densityMclust (mclust v6.1 R package). Input TD segments consists of events meeting the criteria in the TDP classification analysis. For each sample, solutions of one to four possible Gaussian components were used to fit the log10-transformed TD lengths in kilobases. The optimal solution is selected via Bayesian information criteria (default in MClust), and components in the optimal solution with mixed weights ≤ 0.1 or variance. The estimated means for the remaining components determine the size groups: 1-100kb (Group 1), 100kb-1Mb (Group 2), 1-10Mb (Group 3). For solutions with > 1 group (i.e. > 1 component), additional groups were defined: Group 4 (Group 1 + Group 2), Group 5 (Group 1 + Group 3), and Group 6 (Group 2 + Group 3). TD Group size analysis is shown separately for WES and WGS due to differences in copy number segment size resolution – WGS is expected to have higher resolution and therefore smaller TD segments are better represented. Finally, the distribution of the estimated component means is shown for all WES (n=26) and WGS (n=20) samples.

### Genomic regions and genes altered by TDs

Three different sets of genomic regions were analyzed to determine the frequency of alteration by TDs in TDP+ and TDP-samples. Simple TD segment events were defined based on criteria set in the TDP classification analysis*: 1) Tiled 50kb windows across the genome.* The number of TD events overlapping each 50kb window was computed across all TDP+ samples (n=46) and all TDP-samples (n=777) with >1 simple TD event meeting the TDP classification criteria. A two-sided c^2^-test of independence with Bonferroni multiple-test correction was used to determine enrichment of TD alteration between TDP+ and TDP-groups*; 2) COSMIC Cancer Gene Census oncogenes.* The Cancer Gene Census file was from a 1/27/2019 release; gene coordinates in this file were used. Genes that had a ‘Role in Cancer’ value containing the string “oncogene” or “fusion” were considered in this list of 433 oncogenes. A gene was considered to be altered by a TD if the gene coordinates were fully contained within a TD event (i.e. the TD event spans the entire gene). For each gene, a two-sided Fisher’s exact test was used to determine enrichment of TD alteration frequency between TDP+ and TDP-groups. Bonferroni correction was applied*; 3) COSMIC Cancer Gene Census tumor suppressor genes.* The same Cancer Gene Census file was used as above. Genes that had a ‘Role in Cancer’ value containing the string “TSG” or “fusion” were included in this list of 394 genes. A gene was considered to be altered by a TD if either “start” or “end” or both boundaries are located within the gene coordinates (i.e. either or both breakpoints transects the gene). A Fisher’s exact test, followed by Bonferroni correction was applied as above.

Analyses were performed using GRCh38, except for the HMF cohort which was GRCh37 and required liftOver or matching of gene symbols.

### Multi-tumor tandem duplication and phylogenetic analysis of rapid autopsy samples

For UW patient 05-217, input TD segments consisted of events meeting the criteria in the TDP classification analysis. The complete set of TDs within the patient was defined as the union of TD events were taken across all samples within the patient. A matrix consisting of samples by TD event was constructed using 1 for presence of the TD in the sample or 0 if it was absent. A TD event was considered present if the proportion of the overlap by width was ≥ 0.9. This matrix was used to determine the number of overlapping events in pairwise, triplets, and all samples within the patient. This matrix was also used as input into neighbor-joining tree estimation (ape R package) using Manhattan distance to construct a phylogenetic tree with a rooted-tree configuration. The tree branching configuration and general branch length was used to produce a custom representation presented in Figure 2b.

### Annotating gene alteration by copy-number

Copy-number segments were excluded if their cellular fraction was lower than 0.8, except for those which were determined as copy neutral or copy-number greater than 4. The gene annotation was based on known protein coding genes from GenCode release 30 (GRCh38.p12). For each gene, its copy-number was assigned to the copy-number value and LOH status of the segment that has the largest overlap with it. The gene-level copy-number was normalized based on ploidy of the corresponding sample, with autosomal genes normalized by the inferred ploidy rounded to nearest integer, and X-linked genes normalized by half such value. Then the copy-number status of each gene was categorized based on the following criteria: (i) Amplification. Normalized gene-level copy-number is greater than or equal to 2.5; (ii) Gain. Normalized gene-level copy-number is between 1.5 and 2.5; (iii) Homozygous deletion. Normalized gene-level copy-number is 0; (iv) Deletion with LOH. Normalized gene-level copy-number is between 0 and 1, and LOH status was found; (v) Copy neutral LOH. Normalized gene-level copy-number is 1 and LOH status was found.

### Annotating gene alteration by structural variant

Gene coordinates were based on ENSEMBL v33 of hg38. Gene body region of one gene was defined as the widest region of all known isoforms collapsed. Gene flanking region was defined as the corresponding two 1 Mb regions next to the gene body region on 5’-end and 3’-end, respectively. Gene alteration status by genome rearrangements was defined based on the breakpoints and directions of involving structural variant events. A gene in one WGS sample (gene-sample pair) was considered having gene transecting events if any breakpoints of SV events were located within the gene body region. If the gene transecting status did not apply, then this gene-sample pair was examined for gene flanking status if the breakpoints of any intra-chromosomal SV events, including tandem duplications, deletions, and inversions, were located within the gene flanking regions. Additionally, translocation events including intra-chromosomal balanced and unbalanced events which spanned over 10 Mb, and inter-chromosomal translocation events were considered altering the gene flanking regions if any of their breakpoints was in the gene flanking region, and the direction of the SV was going towards the gene body region. The alteration status of rearrangements for each gene-sample pair was exclusive between gene transecting and gene flanking, with the former being prioritized in report.

### Annotating gene allelic alteration status

For CDK12, BRCA1, BRCA2, TP53, CHD1, PTEN, and PABL2, allele status was defined using SNV, Indel, CNV, and SV results, and classified into three categories: Bi-allelic loss (BAL), Mono-allelic loss (MAL), and Intact. An event was only included in gene status calling if it passed all filters applied to the corresponding data type. For a given gene, BAL was defined as two or more events or a homozygous deletion within gene boundaries. MAL was defined as one detected loss-of-function event occurring within gene boundaries, while samples annotated as Intact had no recorded variants. Manual curation was performed to confirm gene statuses, with an emphasis being placed on confirming or amending discordant calls between samples from the same patient.

### Loss of heterozygosity (LOH) analysis

The LOH score was defined as the proportion of the genome affected by LOH events (minor copy number = 0). To compute this value, we summed the genomic distance spanned by all segments reported by TitanCNA to have a minor allele copy number of 0, then divided this value by the genomic distance spanned by all segments reported by TitanCNA. Chromosome arms with LOH events spanning more than 75% of their length were excluded from our analysis, as these events are associated with non-HRD mechanisms.

### Transcript Analyses

Sequencing reads were mapped to the hg38 human genome using STAR.v2.7.3a (74). Gene fusions were mapped and quantitated using STAR-Fusion. AR-V7 quantitation was performed as previously described (75). All subsequent analyses were performed in R. Gene level abundance was quantitated using GenomicAlignments (76). Differential expression between groups was assessed using limma (77) filtered for a minimum expression level using the filterByExpr function with default parameters prior to testing. Genome-wide gene expression results were ranked by their limma statistics and used to conduct Gene Set Enrichment Analysis (GSEA) to determine patterns of pathway activity utilizing the curated pathways from within the MSigDB (78). Single sample enrichment scores were calculated using GSVA (79) with default parameters using genome-wide log2 FPKM values as input. Intronic APA analyses were performed using APAlyzer (v1.2.0) using prebuilt hg38 intronic polyadenylation (IPA) and 3’-most exon regions (49). The read cutoff parameter “CUTreads” was set to 5 for analysis of IPA between groups.

### Cell lines and culture

UWB1.289 were a gift from Dr. Elizabeth Swisher (56) and grown in 50:50 RPMI (Thermo 11875093) and MEGM (Lonza CC-3150) with 3% FBS (Thermo 16000044) and 0.5X Pen/Strep (Thermo 15140122). Skov3 (HTB-77), LNCaP (CRL-1740), C4-2B (CRL-3315), 22Rv1 (CRL-2505), and HEK293T (CRL-3216) were received from ATCC. LuCaP cell lines were generated by resecting the tumor implant, dissociating cells by enzymatic digestion and plating cells in DMEM medium with 10% FBS or with various additives used in organoid medium (80). Skov3, LNCaP, C4-2B, and 22Rv1 were grown in RPMI with 10% FBS and 0.5X Pen/Strep. LuCaP 35_CL, LuCaP 189.4_CL, and HEK293T were grown in DMEM (Thermo 11965092) with 10% FBS, 1X GlutaMAX (Thermo 35050061), 1mM additional sodium pyruvate (Thermo 11360070), and 0.5X Pen/Strep. Organoids were grown in customized media as previously described (81). Cells were grown at 37C with 5% CO2. Cell lines underwent DNA fingerprint (STR) confirmation and routine mycoplasma testing (R&D Systems CUL001B) via Fred Hutch Research Cell Bank Services.

### Lentivirus production and transduction

HEK293T cells were seeded at 18 million per T75 flask and transfected the next day with 3μg pMISSION-VSVG, 6μg pMISSION-Gag-Pol, and 9μg transfer plasmid with 36μL of TransIT Leti reagent (Mirus MIR 6604). Cells were changed to fresh media the next day and viral media was collected on day 3 for concentration with Lenti-X concentrator (Takara 631232). Cells were transduced with 8μg/mL polybrene. Viral titer was determined by flow cytometry (for fluorescent vectors) or antibiotic selection titer with antibiotic started 48h after infection and viability assayed 6 days post transduction. For growth assay, cells were transduced at a target multiplicity of infection of 2.

### Vectors and engineered lines

Tet-inducible shRNA vectors were cloned as previously described(82) into EZ-Tet-pLKO-Hygro (Addgene 85972). We generated two customized sgRNA construct backbones: pLenti-sgStuffer-GFP-Puro (Addgene 208349) and pLenti-sgStuffer-mCherry-Puro (Addgene 208350). 22Rv1 CDK12-KO lines were generated by stable lentiviral transduction with two different sgRNA vectors: pLenti-CDK12(sg1)-GFP-Puro and pLenti-CDK12(sg2)-GFP-Puro. The stable CDK12 sgRNA lines were then transduced with pLenti-Cas9.mCherry (Addgene 208342) and single cells were sorted by flow cytometer (Sony SH800S) into 96 well plates. Colonies were expanded and screened by western blot while mutations were confirmed by PCR, Sanger sequencing, and RNA-seq. Skov3 CDK12-KO1 was generated similarly, except that the sgRNA and Cas9 plasmids were transiently transfected before flow sorting for clonal expansion. The Skov3-CDK12-KO1 line has pre-existing puromycin resistance from previous transduction with a non-functional (frame-shifted) lentiviral vector. LuCaP189.4_CL were transduced with FUCGW(83) or FUCGW-CDK12 and flow enriched for GFP to create isogenic pools LuCaP189.4-vec and LuCaP189.4-CDK12. FUCGW was a gift from Dr. John K. Lee, which was modified into FUCGW-DEST (Addgene 208408) and FUCRW-DEST (Addgene 208409) by inserting an attR1-CmR-ccdB-attR2 cassette. CDK12 CDS was amplified with Phusion HF (Thermo F530) and TOPO cloned into the pCR8 entry vector (Thermo K250020) to make pCR8-CDK12 (Addgene 208347). Silent sgRNA-resistant (sgR) mutations were made via assembly cloning (NEB E2621) to generate pCR8-CDK12(sgR) (Addgene 208346), which was then recombined with LR clonase II (Thermo 11791020) into FUCGW-DEST to generate FUCGW-CDK12(sgR). All plasmid sequences were confirmed with full plasmid sequencing (Plasmidsaurus, Eugene, OR). For growth assays, a dual sgRNA vector system was made based on lentiCRISPRv2-Blast (Addgene 83480), generating pLCV2-AAVS1(hU6-sg1-mU6-sg2)-Blast (Addgene 208343), pLCV2-CDK12(hU6-sg1-mU6-sg2)-Blast (Addgene 208344), and pLCV2-CDK13(hU6-sg1-mU6-sg2)-Blast (Addgene 208348). Additional information on the cloning strategy, including the dual sgRNA cloning protocol, is available on the vector pages on addgene.org. Primer. shRNA, and sgRNA sequences are listed in **Table S4**.

### Immunoblot

Lysates were prepared in RIPA buffer (150mM NaCl, 5mM EDTA, 50mM Tris, pH 8.0, 1% IGEPAL CA-630, 0.5% sodium deoxycholate, 0.1% SDS), passed through a 20G needle, pelleted to remove debris, then quantified by BCA assay (Thermo 23225). Loading samples were prepared in LDS sample buffer (Thermo NP0007) with 5% beta-mercaptoethanol and heated at 99C for 10m for denaturation. Samples were run on NuPAGE 4-12% bis-tris gels (Thermo NP0335BOX) with MOPS run buffer (Thermo NP000102) or 3-8% tris-acetate gels (Thermo EA0378BOX) with tris acetate SDS run buffer (Thermo LA0041). Gels were run and transferred in an XCell SureLock module (Thermo EI0002). Wet transfers were performed with NuPAGE transfer buffer (Thermo NP00061) containing 10% methanol for 3h at 30V fixed voltage in a cold room. Additional reagents include BLUEstain2 protein ladder (GoldBio P008-500) and low fluorescence PVDF (BioRad 1620260) and StartingBlock buffer (Thermo 37542). Blots were imaged on a ChemiDoc MP system (BioRad 12003154) with ImageLab software (v6.1). Antibody information is in **Table S4**.

### RAD51+γH2A.X immunofluorescence and foci quantification

Cells were seeded on 12mm circle coverslips coated with poly-d-lysine (50ug/mL PBS, 400uL per 12-well, 1h at 37C) (Thermo A3890401). Cells were placed in a cesium-137 irradiator for a 6 Gy exposure or mock treated then returned to the tissue culture incubator and fixed 3h post irradiation with 4% paraformaldehyde/PBS (Thermo AAJ19943K2) for 10m. Fixed cells were then quenched with 125mM glycine/500mM tris pH 7.4 for 10m and permeabilized with 0.5% Triton X-100 in TBS (50mM tris, 150mM sodium chloride, pH 7.4) for 5m then washed once with TBS/T (TBS with 0.1% Tween 20). Next, cells were blocked with Dual Endogenous Enzyme Buffer (Agilent S200389-2) for 10m at room temp then washed twice with TBS/T. Cells were incubated overnight at 4C with RAD51 (Sigma ZRB1492) and γH2A.X (Sigma 05-636) primary antibodies in 1% BSA(TBS/T) (**Table S4**). After primary incubation, cells underwent two washes in TBS/T and were incubated with HRP-anti-rabbit secondary (Leica PV6119) for 30m at room temp. Next, cells were washed three times with TBS/T and then incubated for tyramide amplification and Alexa 568 staining according to the vendor protocol (Thermo B40956). Cells were washed three times with TBS/T and then incubated 1h at room temp with Alexa Fluor 488 anti-mouse (Thermo A32723) or Alexa Fluor 633 anti-mouse (Thermo A-21052) for 22Rv1 lines. Following a TBS/T wash, cells were stained with DAPI (2ug/mL in TBS/T) for 10m at room temp, washed twice in TBS/T and once in water, then mounted onto glass slides with ProLong Gold (Thermo P36930). Slides were imaged on a Zeiss LSM800 confocal microscope running Zen software (v2.3) with a 20X/0.8NA objective, 40μm pinhole, and three 1.5μm z-steps. Image stacks were flattened with a maximum intensity projection and foci were quantified using imageJ (version 1.53q)(84) to outline nuclei and count foci using the FindMaxima tool. Data were plotted using GraphPad PRISM (v10.0.1) showing number of foci per nucleus with a line at the mean with at least 150 nuclei per group. Statistical significance was determined by ANOVA (Kruskal-Wallis).

### Quantitative real time PCR (qPCR)

RNA was extracted with Qiagen RNeasy kit (Qiagen 74004) and quantified by Qubit (Thermo Q10211). Reverse transcription was performed with ProtoScript II (NEB M0368) using 4μM poly-dT plus 1μM random hexamer primers. qPCR was performed in 384-well plates (15μL reactions) with Power SYRB Green Mix (Thermo 4367660) and run on a QuantStudio 6 Flex (Thermo 4485691). Relative expression was standardized (ΔCT) to zero-centered average of four housekeeping primer sets (*18S*, *RPL19*, *ACTB*, *GAPDH*) and normalized (ΔΔCT) as described in each experiment, generating log2(fold change). Primer information is in **Table S4**.

### Drug dose response curves

Cells were seeded at relatively low densities in 96-well plates (500-1,000 for Skov3 and UWB1.289; 1,000-4,000 for 22Rv1 and LNCaP; 10,000-20,000 for LuCaP189.4_CL).

Cells were seeded 24-48h before adding drug. There were no media changes until harvest unless otherwise noted. Cells were kept at room temp for 30m after seeding before transferring to incubator and were kept in a humidified chamber (e.g. plastic box with water reservoir) to promote even seeding and reduce edge effects.(85) Cells were harvested by adding 50uL of CellTiterGlo 2.0 reagent (Promega G9243) per well, incubating 20 minutes, and reading luminescence on a BioTek plate reader with Gen5 software. For organoid experiments, 5,000-10,000 cells were seeded per 15uL Matrigel droplet (Corning 354234) (2:1 gel:cell suspension ratio) in 96-well round bottom plates. At harvest, 50uL of CellTiterGlo was added per well, incubated 20m, then wells were mixed by pipette and transferred to solid black plates before reading luminescence. Data were normalized vs mean of media-only wells and plotted with PRISM showing mean-/+stdev and polynomial curve fitting (inhibitor vs response, variable slope, four parameter) for effective concentration 50 (EC50) calculation for concentration of drug at 50% relative viability.

### sgRNA growth assays

GFP-tagged cells (transduced with FUCGW) were infected with sgRNA lentivirus in 6-well plates. 24-48h later 10ug/mL blasticidin was added for selection and on day 5 post-infection cells were seeded into 384-well plates (Corning 353962) which were read once per 24h on a Cytation5 (BioTek) with BioSpa using the GFP channel and 1.25X objective. GFP confluence was calculated in ImageJ setting a threshold and counting GFP positive pixels. LuCaP189.4_CL cells were seeded on low adherence plates (Perkin Elmer 6057800) and quantified with Gen5 software.

### PDX tumor growth experiment

LuCaP PDX lines were implanted subcutaneously into male NSG mice (NOD scid gamma) (one tumor per mouse). Treatment began when tumors reached 150mm^3 volume as measured by calipers. SR4835 was purchased from SelleckChem (S8894). Fresh solutions were prepared by first solubilizing drug in DMSO (Sigma D2650) and then diluting the stock dropwise 1:10 into 30% cyclodextrin (w/v) in water (Sigma H107). Mice were given 20mg/kg SR4835 or vehicle by oral gavage 5 days on / 2 days off.(21) Tumor volume and mouse weights were measured 3 times/week using calipers, with volume calculated as 4/3*π*L*W*H/8. Animal use followed Fred Hutch IACUC approved protocol (#51077). Mice were sacrificed at 28 days treatment, maximum allowed tumor size (tumor avg. diameter >2cm), or if deemed necessary by comparative medicine staff. Tumors were excised and weighed upon sacrifice.

## CODE AVAILABILITY

Analysis code for custom analysis and pipeline configuration settings are accessible at https://github.com/GavinHaLab/CDK12-CRPC-paper.

## ACKNOWLEDGEMENTS

We would like to thank the patients who generously donated tissue that made this research possible. We would also like to thank Lisa Ang and Talina Nunez for assistance with mouse studies and Pushpa Itagi for assistance with genomic analyses. We thank Evan Yu, Heather Cheng, Jessica Hawley, Michael Schweizer, Lawrence True, Meagan Chambers, Martine Roudier, and the rapid autopsy teams for their contributions to the University of Washington Medical Center Prostate Cancer Donor Rapid Autopsy Program and the development of the LuCaP PDX models. We thank Arul Chinnaiyan and Dan Robinson for generating and sharing CDK12-mutant tumor transcriptome profiles. This research was supported by the Genomics and Bioinformatics core and Comparative Medicine Shared Resource, RRID:SCR_022606 and RRID:SCR_022610, of the Fred Hutch/University of Washington/Seattle Children’s Cancer Consortium, P30 CA015704. This work was also supported by awards from the CDMRP PC200262, PC200262P1, PC180295, the Pacific Northwest Prostate Cancer SPORE P50CA97186, PO1CA163227, R21CA277368, F32CA243286, R50 CA274336-02, DP2CA280624, R01CA280056, the Institute for Prostate Cancer Research and the Prostate Cancer Foundation.

## FIGURE LEGENDS

**Supplemental Figure S1.**
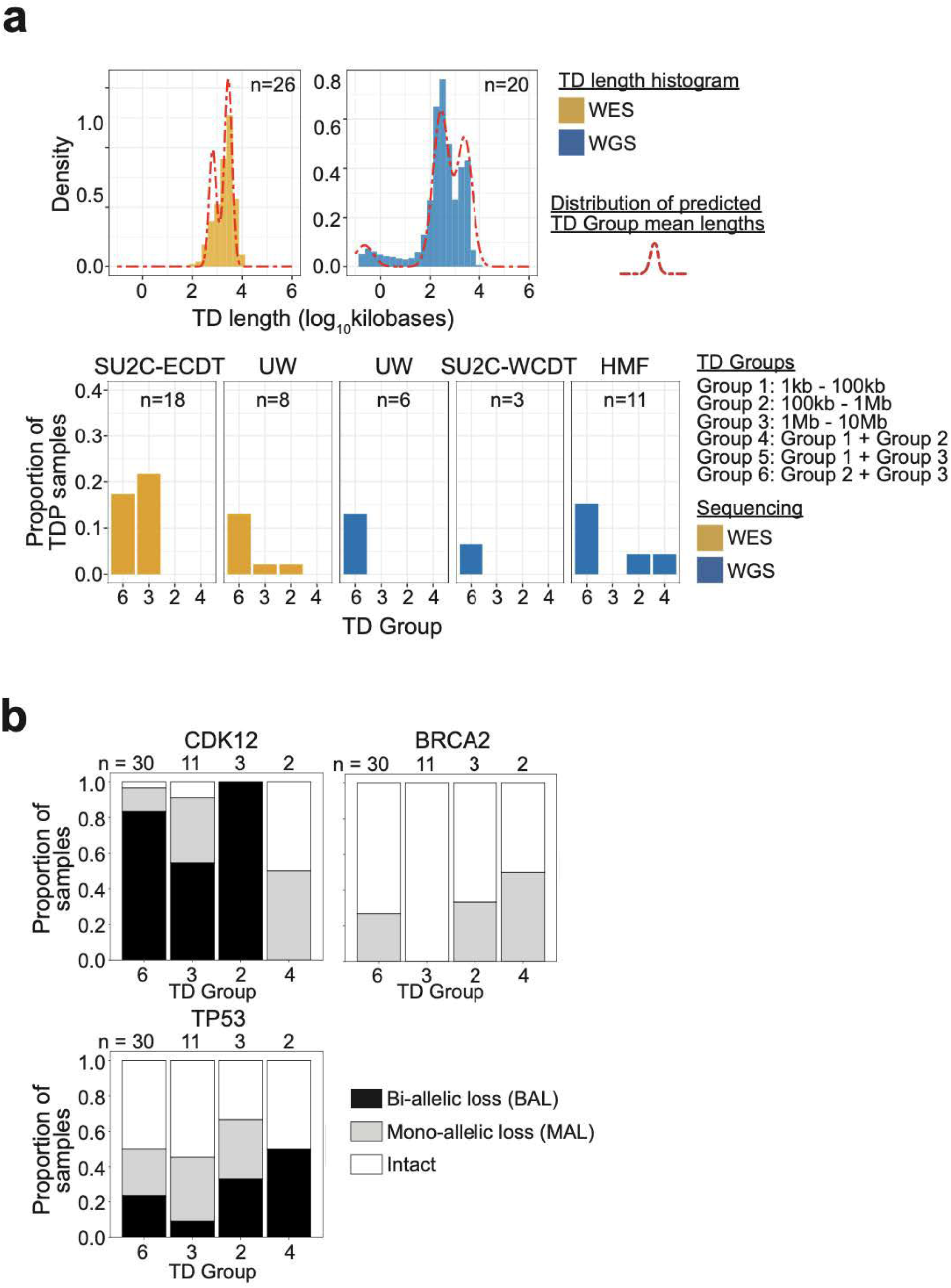
CDK12 genomic alterations and tandem duplication phenotypes. **(a)** Tandem duplication (TD) size analysis confirms multiple modes of TD lengths (top) in whole exome (WES; orange) and whole genome sequencing (WGS; blue) data for 46 tumors with a TDP. Fitting mixtures of Gaussian distributions to each of 26 WES and 20 WGS samples provided estimates of Gaussian means. The distribution of these mean values across 26 WES cases and 20 WGS cases are shown in red dotted lines; samples may have multiple mean values, one for each estimated mixture. The mixture means are categorized into six possible TD Groups based on the length ranges (bottom). The proportion of each TD Group are shown for the four CRPC cohorts. **(b)** The proportions of *CDK12*, *BRCA2*, and *TP53* genomic alteration status, including mono-allelic and bi-allelic losses and intact, for the 46 TDP cases compared across TD Groups.

**Supplemental Figure S2.**
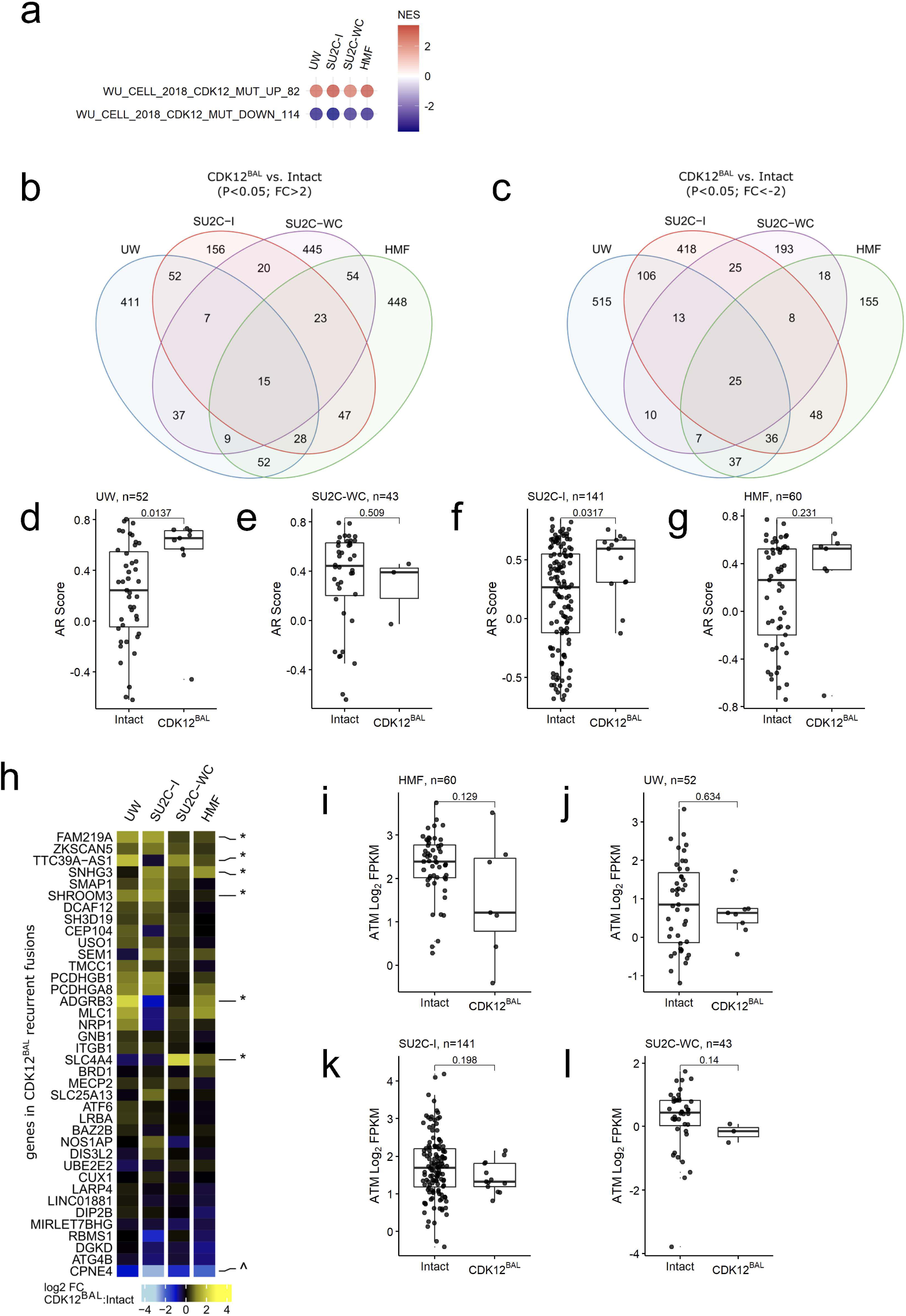
Gene expression alterations in *CDK12^BAL^* prostate cancers. (**a**) GSEA of CDK12-mut up and down signatures reported in Wu et al., 2018 applied to mCRPC cohorts with versus without *CDK12^BAL^*. (Normalized enrichment scores (NES) shown in heatmap on plot, all enrichment false discovery rates (FDR) <0.0001). **(b-c)** Overlap of differentially expressed genes up-regulated (**c**) or down-regulated **(d)** across mCRPC cohorts with versus without *CDK12^BAL^* (significance level shown on plot.) **(d-g)** RNAseq based quantitation of AR pathway activity in mCRPC cohorts with versus without *CDK12^BAL^* (GSVA scores; Wilcoxon rank test p-values shown). **(h)** Differential expression of genes contained within *CDK12^BAL^*-specific and recurrent fusions. * = Up-regulated or ^ down-regulated with p<0.05, FC>abs(2) in at least 1 cohort. **(i-l)** RNAseq based transcript abundance levels of ATM in mCRPC cohorts with versus without *CDK12^BAL^* (Wilcoxon rank test p-values shown).

**Supplemental Figure S3.**
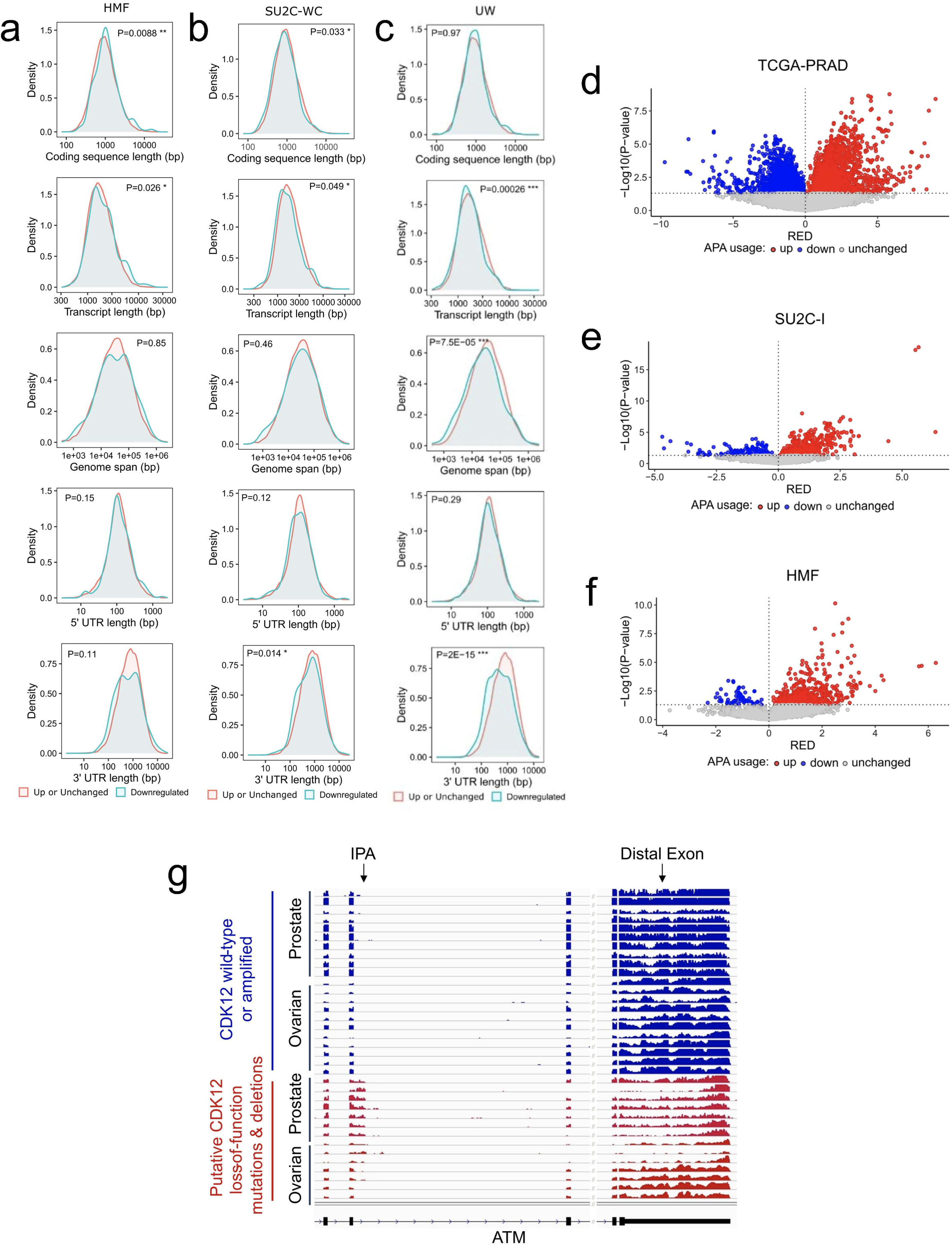
APA usage in *CDK12^BAL^* prostate cancers. **(a-c)** Association of increased/unchanged or decreased (p<0.05; FC<-2) transcript abundance levels in relation to gene size features in tumors without or with CDK12^BAL^ in the HMF, SU2C-WC and UW mCRPC cohorts. **(d-f)** APAlyzer analysis for up– and down-regulated APA usage using RNA-seq from mCRPC data sets comparing CDK12^BAL^ cases vs CDK12(intact) controls: (c) TCGA-PRAD, (d) SU2C-I, and (e) HMF. RED = relative expression difference; each point is a different APA site. **(g)** Transcript pile-up of reads mapping to exons demonstrating increased transcript reads corresponding to an IPA in the ATM gene in TCGA prostate (PRAD) and ovarian primary tumors with CDK12 alterations versus tumors with intact CDK12 and diminished transcripts mapping to the distal 3’ exon.

**Supplemental Figure S4.**
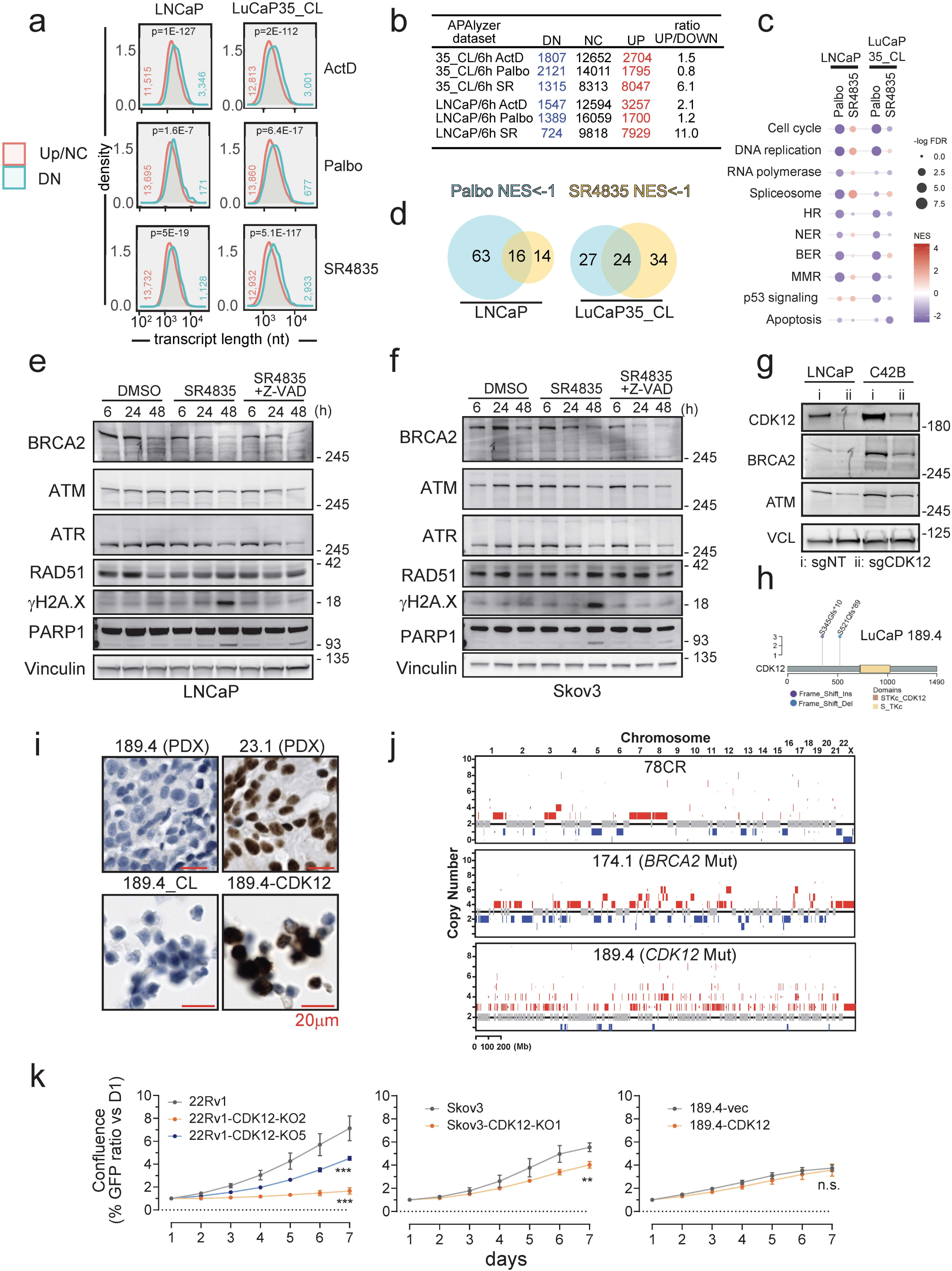
Effects of acute CDK12 inhibition. **(a)** Treatment with ActD, Palbo, and SR4835 all lead to downregulation of longer transcripts. Distribution by transcript length (nucleotides/nt) of downregulated genes by RNA-seq (<-2 fold, FDR <0.05, ‘n’ depicted on plot) in LNCaP and LuCaP35_CL prostate cancer cell lines following six hours of exposure to vehicle (DMSO), CDK4/6 inhibitor palbociclib (Palbo, 10uM), broad RNA Pol-II inhibitor actinomycin D (ActD, 5ug/mL), or CDK12/13 inhibitor SR4835 (200nM) (n=3). Plots were made with ShinyGO 0.80 (76) and show significance (t-test) of downregulated vs upregulated or unchanged genes. **(b)** CDK12 inhibition increases the number and ratio of upregulated APAs. Table with the number of APA sites down (DN), no change (NC), or up (UP) upon treatment vs vehicle and the ratio UP/DOWN, indicating any skew towards the IPA phenotype. **(c-d)** Cell arrest and CDK12 inhibition both lead to HR pathway downregulation. **(c)** Selected KEGG pathway enrichment for DNA-repair related pathways focusing on changes from general G1/S arrest (palbociclib) vs CDK12 inhibition (SR4835). (d) Many SR4835 downregulated pathways overlap with cell cycle arrest. Venn diagram showing high overlap of KEGG pathways downregulated (NES <-1) with acute palbociclib or SR4835 treatment (6h). **(e-f)** Effects of pharmacological CDK12/13 inhibition. Similar experiments as in Fig. 4f using LNCaP (e) and Skov3 (f) cells showing the effect on DNA repair and apoptotic proteins with SR4835 treatment with or without Z-VAD caspase inhibitor. **(g)** LNCaP and C42B were transduced with lentivirus containing dual sgRNAs against CDK12 or nontargeting controls. Lysates were harvested 7 days post infection and analyzed by immunoblot. **(h)** LuCaP189.4 carries bi-allelic loss of function *CDK12* mutations. Diagram showing the two frameshift mutations carried by the LuCaP189.4 PDX upstream of the key functional kinase domain (yellow). **(i)** LuCaP 189.4 does not express CDK12 protein. Immunohistochemical (IHC) staining (with amplification) for CDK12 on FFPE sections from LuCaP PDX tumors or cell spots from LuCaP189.4_CL and LuCaP189.4-CDK12 cell lines. Counterstained with hematoxylin. **(j)** LuCaP189.4 shows hallmark TDP genomic pattern. Copy number plot from exome-seq of three LuCaP PDX lines (78CR, 174.1, and 189.4). **(k)** *CDK12* knockout lines grow slower than parental lines. GFP tagged cells were grown for seven days and monitored by GFP imaging. Graphs show GFP confluence normalized to day 1 for each line with mean-/+stdev (n=5). Significance vs parental line was determined by two way ANOVA.

**Supplemental Figure S5.**
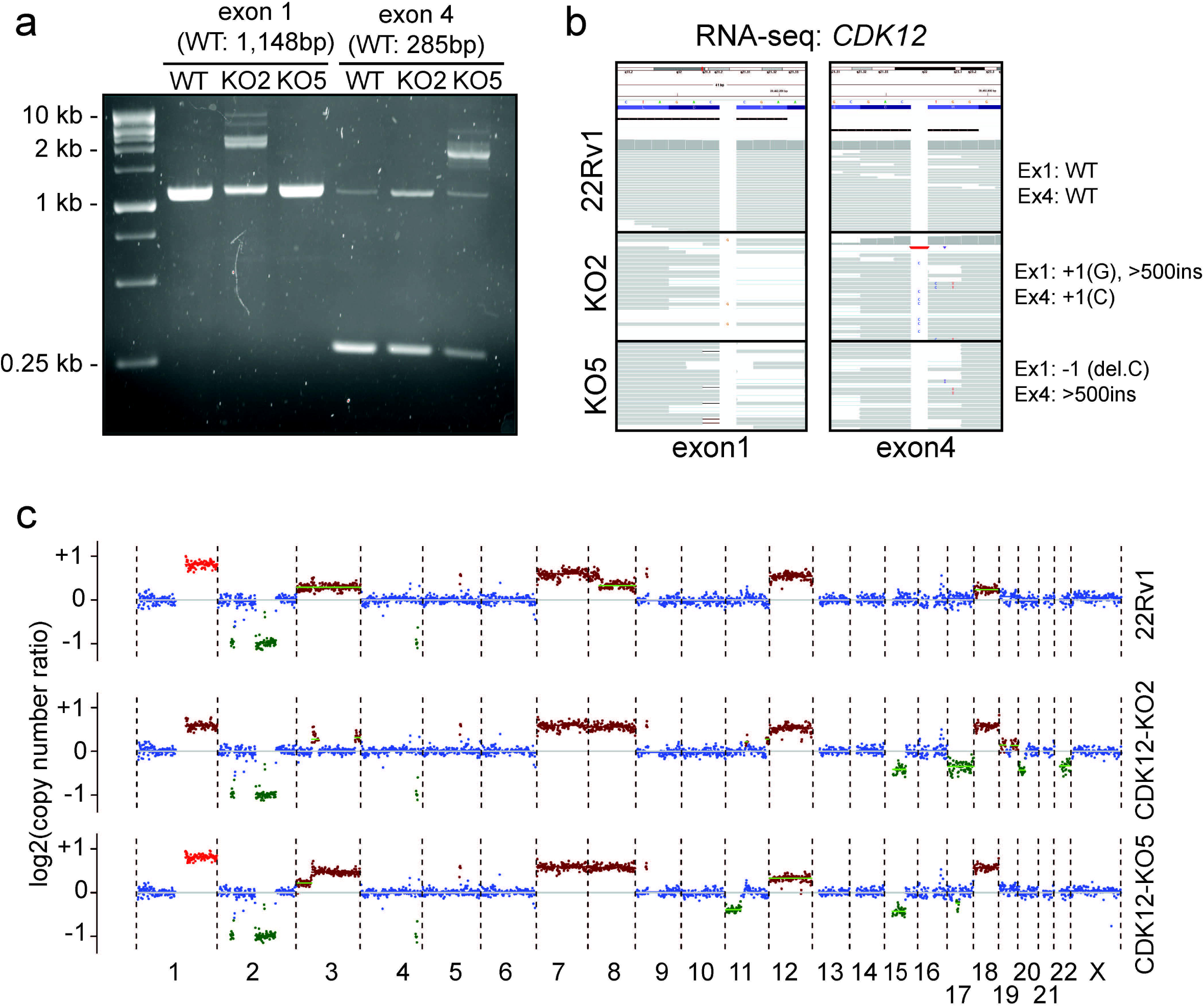
Validation of 22Rv1 CDK12 KO lines. **(a)** PCR was performed on genomic DNA from 22Rv1 lines to amplify the sgRNA targeted sites in *CDK12* exon1 and exon4. Note the large products in KO2 (exon1) and KO5 (exon 4) indicating large genomic insertion events. **(b)** RNA-seq reads from the 22Rv1 lines show presence of frameshift indels in the CRISPR clones. **(c)** Low coverage WGS was performed on the clones and plotted for copy number alterations, with no obvious sign of a TDP pattern (as can be seen in Fig. 2a and S4g).

**Supplemental Figure S6.**
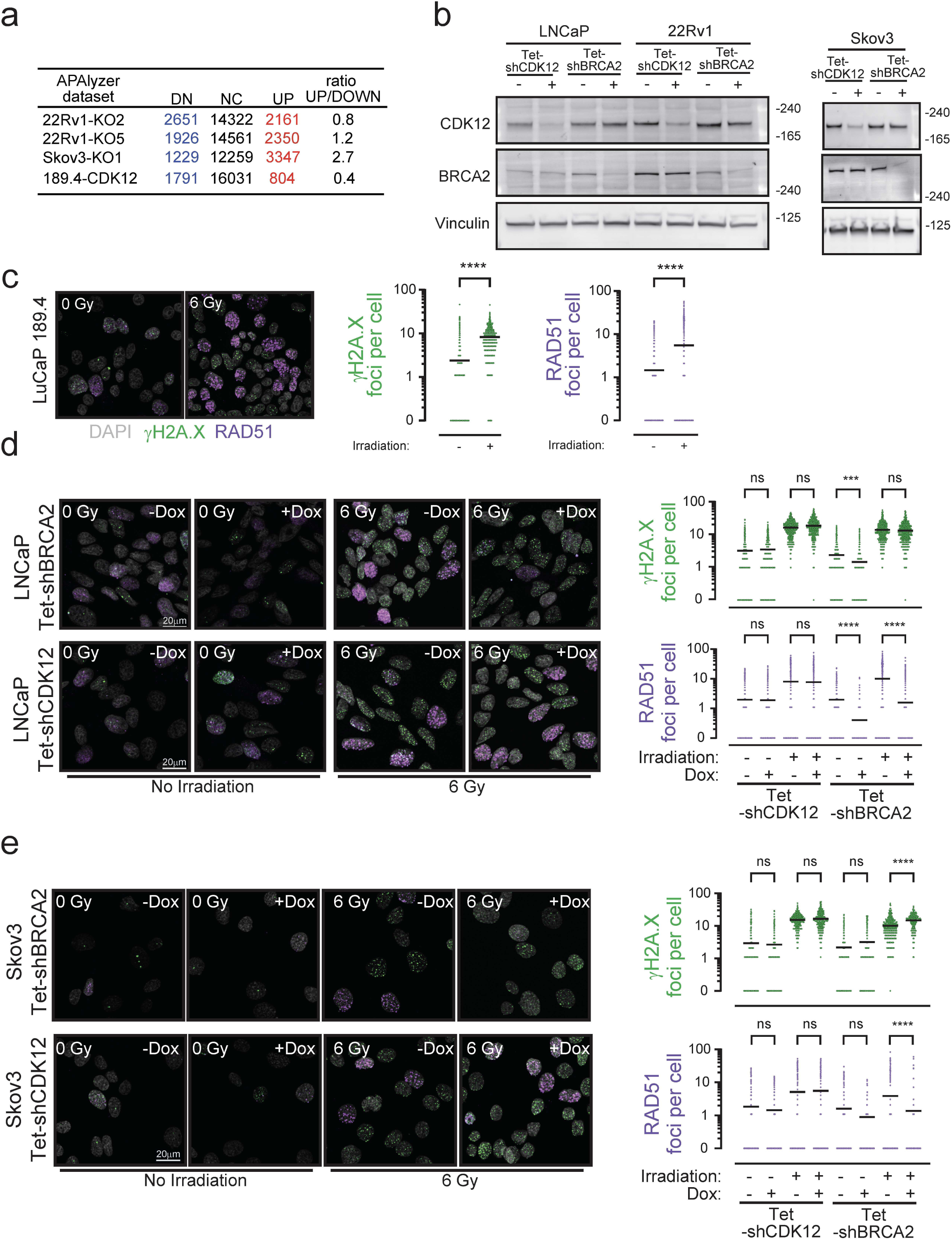
Assessments of HR competency in cells with CDK12 loss. **(a)** Stable CDK12(−) cells show fewer upregulated intronic APAs. Table with the number of APA sites down (DN), no change (NC), or up (UP) in isogenic paired models (CRISPR KO clones vs parental, or 189.4-CDK12 vs 189.4-vec). The ratio UP/DOWN indicates skew towards the IPA phenotype. **(b)** Validation of Tet-shRNA lines. Western blot with lysates from Tet-shRNA lines treated four days −/+ 100ng/mL doxycycline. **(c)** LuCaP189.4_CL cells are RAD51 competent. Irradiation and immunostaining (same as in Fig 5f). Cells were exposed to 6Gy IR and fixed at 3h. Immunofluorescence staining was performed for γH2A.X and RAD51 and images were acquired by confocal microscopy. Left: representative images (white: DAPI, green: γH2A.X, purple: RAD51). Right: quantification of images (∼200-500 cells analyzed per treatment). Line is at mean and significance was determined by unpaired t-test (Mann-Whitney). **(d-e)** *CDK12* knockdown does not prevent RAD51 foci. Additional immunostaining (same as in Fig 5f) using LNCaP (d) and Skov3 (e) with Tet-shCDK12 or Tet-shBRCA2. Cells were treated four days −/+ dox. Graphs show mean with significance determined by one-way ANOVA (Kruskal-Wallis).

**Supplemental Figure S7.**
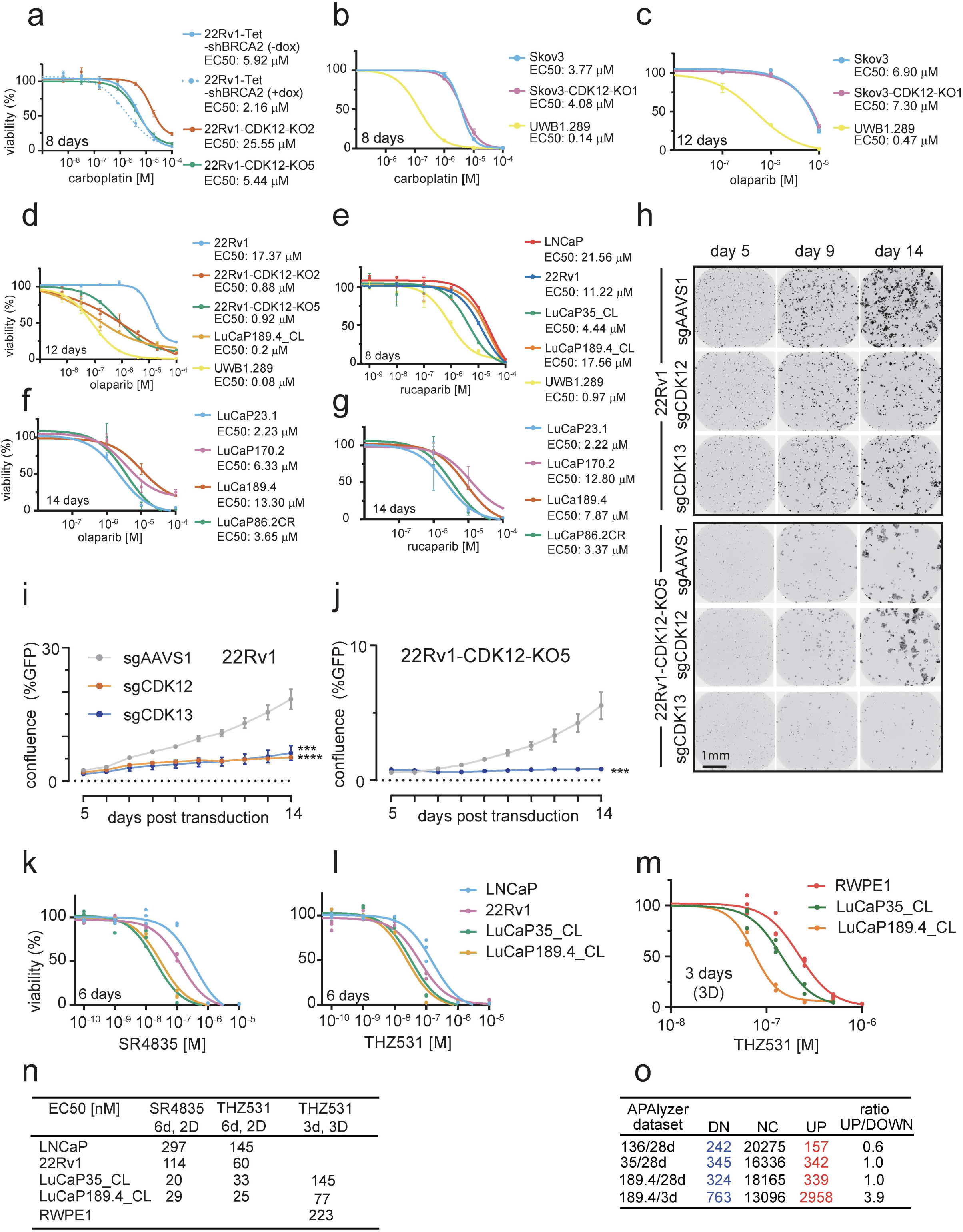
Drug sensitivities in prostate cancers with CDK12 loss. **(a-b)** CDK12 loss does not confer HRd-expected platinum or PARPi sensitivity. Dose response curves for prostate cancer (a) or ovarian cancer (b) cell lines treated 8 days with carboplatin (n=3 for prostate lines, n=4 for ovarian lines). EC50 values are shown on the legend. **(c)** Ovarian cancer cells (n=4) were treated 12 days with olaparib. **(d)** Prostate cancer lines and UWB1.289 (n=3) were treated 12 days with olaparib. **(e)** Prostate cancer lines and UWB1.289 (n=3) were treated 8 days with rucaparib. **(f-g)** LuCaP189.4 organoids do not show obvious PARPi sensitivity. LuCaP PDX tumors were dissociated into organoids and treated (at passage 3) with olaparib (f) or rucaparib (g) for 14 days. Plots show mean-/+stdev (n=4). **(h-j)** CDK13 sgRNA is detrimental in 22Rv1 lacking *CDK12*. GFP tagged 22Rv1 or 22Rv1-CDK12-KO5 were transduced with sgRNAs and monitored by imaging. Example images are in (h). Plots show growth rates for 22Rv1 (i) and 22Rv1-CDK12-KO5 (j) by confluence (%GFP-/+stdev, n=5) with significance vs sgAAVS1 determined by two-way ANOVA. **(k-n)** LuCaP189.4_CL shows sensitivity to CDK13 inhibition. Dose response curves from four prostate cancer lines, including CDK12^BAL^ LuCaP189.4_CL, treated six days with the CDK12/13 inhibitors SR4835 (k) or THZ531 (l). 3D cultured spheroids were treated three days with THZ531, confirming LuCaP189.4 heightened sensitivity (m). Calculated EC50 values (nM) are listed in table (n). **(o)** APAlyzer analysis from SR4835 treated PDX tumors. Three tumors of each group from Fig. 6j plus three day treated LuCaP189.4 were analyzed by RNA-seq. The table shows the number of intronic APA sites down (DN), no change (NC), or up (UP) in treated vs vehicle comparisons. The ratio UP/DOWN indicates skew towards the IPA phenotype.

**Supplemental Figure S8.**
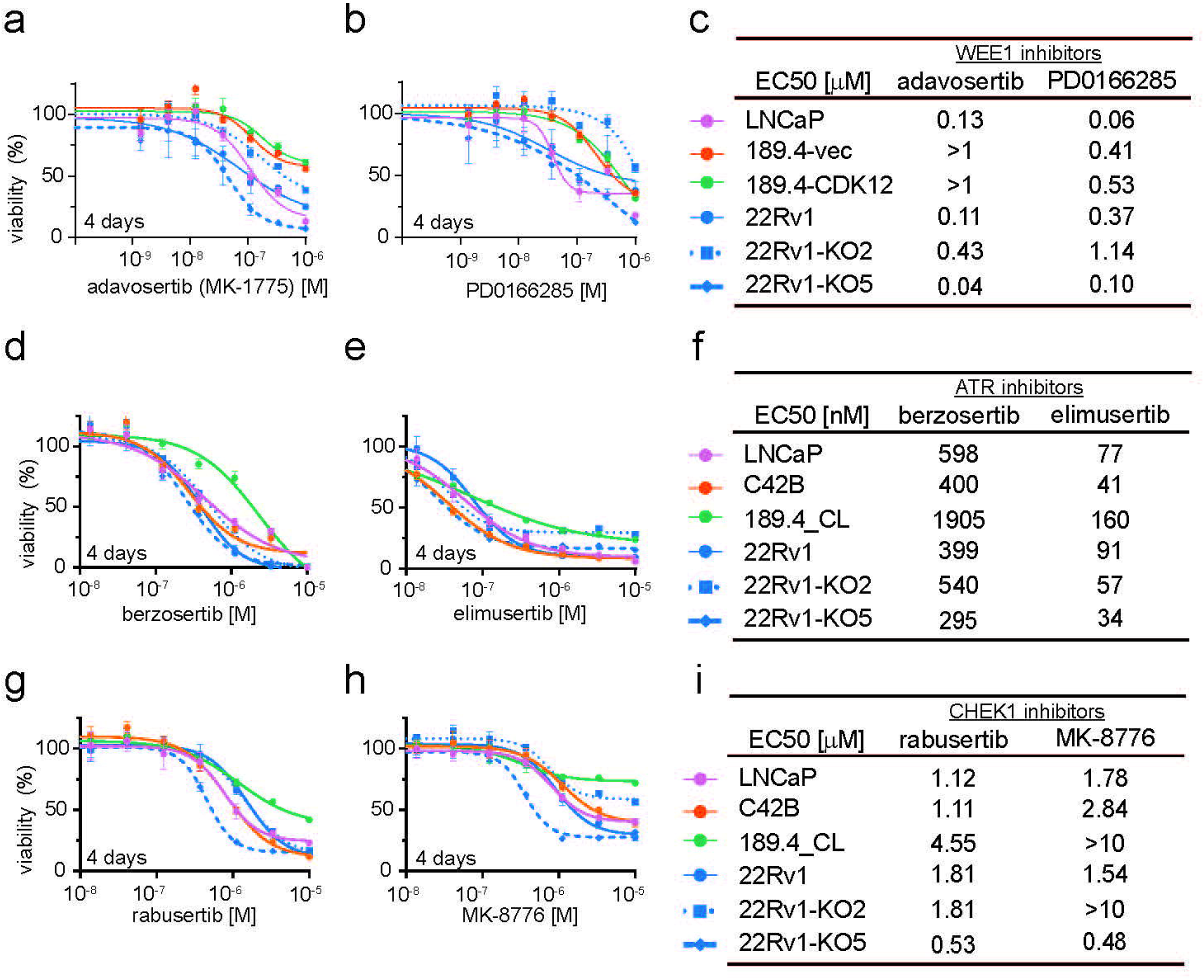
Effects of WEE1, ATR and CHEK1 inhibitors toward prostate cancers with CDK12 loss. **(a-c)** One CDK12-KO prostate line, 22Rv1-CDK12-KO5, shows increased sensitivity to WEE1 inhibition. Prostate cancer lines were treated with WEE1 inhibitors adavosertib/MK-1775 (a) and PD0166285 (b) for four days. Dose response curves show relative viability vs drug concentration. Plots show mean-/+stdev (n=4). Legend and EC50 values (μM) are shown in (c). **(d-f)** CDK12-KO prostate lines do not show sensitivity to ATR inhibitors. Prostate cancer lines were treated with ATR inhibitors berzosertib/VX-970 (d) and elimusertib/BAY-1895344 (e) for four days. Dose response curves show relative viability vs drug concentration. Plots show mean-/+stdev (n=4). Legend and EC50 values (nM) are shown in (f). (**g-i**) One CDK12-KO prostate line, 22Rv1-CDK12-KO5, shows increased sensitivity to CHEK1 inhibition. Prostate cancer lines were treated with CHEK1 inhibitors rabusertib (g) and MK-8776 (h) for four days. Dose response curves show relative viability vs drug concentration. Plots show mean-/+stdev (n=4). Legend and EC50 values (μM) are shown in (i).

